# Jurassic NLR: conserved and dynamic evolutionary features of the atypically ancient immune receptor ZAR1

**DOI:** 10.1101/2020.10.12.333484

**Authors:** Hiroaki Adachi, Toshiyuki Sakai, Jiorgos Kourelis, Hsuan Pai, Jose L. Gonzalez Hernandez, Abbas Maqbool, Sophien Kamoun

**Affiliations:** The Sainsbury Laboratory, University of East Anglia, Norwich Research Park, NR4 7UH, Norwich, UK; Graduate School of Biological Sciences, Nara Institute of Science and Technology, Ikoma 630-0192, Japan; Laboratory of Crop Evolution, Graduate School of Agriculture, Kyoto University, Mozume, Muko, Kyoto 617-0001, Japan; Agronomy, Horticulture and Plant Sciences Department, South Dakota State University, Brookings, South Dakota, US

## Abstract

In plants, NLR immune receptors generally exhibit hallmarks of rapid evolution even at the intraspecific level. We used iterative sequence similarity searches coupled with phylogenetic analyses to reconstruct the evolutionary history of ZAR1, an atypically conserved NLR that traces its origin to early flowering plant lineages ∼220 to 150 million years ago (Jurassic period). We discovered 120 ZAR1 orthologs in 88 species, including the monocot *Colacasia esculenta*, the magnoliid *Cinnamomum micranthum* and the majority of eudicots, notably the early diverging eudicot species *Aquilegia coerulea*. Ortholog sequence analyses revealed highly conserved features of ZAR1, including regions for pathogen effector recognition, intramolecular interactions and cell death activation. We functionally reconstructed the cell death activity of ZAR1 and its partner receptor-like cytoplasmic kinase (RLCK) from distantly related plant species, experimentally validating the hypothesis that ZAR1 has evolved to be a partner with RLCKs early in its evolution. In addition, ZAR1 acquired novel features, such as a C-terminal integration of a thioredoxin-like domain. ZAR1 duplicated into two paralog families, which underwent distinct evolutionary paths. We conclude that ZAR1 stands out among angiosperm NLRs for having experienced relatively limited gene duplication and expansion throughout its deep evolutionary history. Nonetheless, ZAR1 did also give rise to non-canonical NLR proteins with integrated domains and degenerated molecular features.

## INTRODUCTION

Plants immune receptors, often encoded by disease resistance (*R*) genes, detect invading pathogens and activate innate immune responses that can limit infection (Jones and Dangl, 2006). A major class of immune receptors is formed by intracellular proteins of the nucleotide-binding leucine-rich repeat (NLR) family (Dodds and Rathjen, 2010; Jones et al., 2016; Kourelis and van der Hoorn, 2018). NLRs detect host-translocated pathogen effectors either by directly binding them or indirectly via host proteins known as guardees or decoys. NLRs are arguably the most diverse protein family in flowering plants (angiosperms) with many species having large (>100) and diverse repertoires of NLRs in their genomes (Shao et al., 2016; Baggs et al., 2017; Kourelis et al., 2021). They typically exhibit hallmarks of rapid evolution even at the intraspecific level (Van de Weyer et al., 2019; Lee and Chae, 2020; Prigozhin and Krasileva, 2020). Towards the end of the 20^th^ century, Michelmore and Meyers (1998) proposed that NLRs evolve primarily through the birth-and-death process. In this model, new NLRs emerge by recurrent cycles of gene duplication and loss—some genes are maintained in the genome acquiring new pathogen detection specificities, whereas others are deleted or become non-functional through the accumulation of deleterious mutations. Such dynamic patterns of evolution enable the NLR immune system to keep up with fast-evolving effector repertoires of pathogenic microbes. However, as already noted over 20 years ago by Michelmore and Meyers (1998), a subset of NLR proteins are slow evolving and have remained fairly conserved throughout evolutionary time (Wu et al., 2017; Stam et al., 2019). These “high-fidelity” NLRs (per Lee and Chae, 2020) offer unique opportunities for comparative analyses, providing a molecular evolution framework to reconstruct key transitions and reveal functionally critical biochemical features (Delaux et al., 2019). Nonetheless, comprehensive evolutionary reconstructions of conserved NLR proteins remain limited despite the availability of a large number of plant genomes across the breadth of plant phylogeny. One of the reasons is that the great majority of NLRs lack clear-cut orthologs across divergent plant taxa. Here, we address this gap in knowledge by investigating the macroevolution of ZAR1 (HOPZ-ACTIVATED RESISTANCE1), an atypically ancient NLR, and asking fundamental questions about the conservation and diversification of this immune receptor throughout its deep evolutionary history.

NLRs occur across all kingdoms of life and generally function in non-self perception and innate immunity (Jones et al., 2016; Uehling et al., 2017). In the broadest biochemical definition, plant NLRs share a multidomain architecture typically consisting of a NB-ARC (nucleotide-binding domain shared with APAF-1, various R-proteins and CED-4) followed by a leucine-rich repeat (LRR) domain. Angiosperm NLRs form several major monophyletic groups with distinct N-terminal domain fusions (Shao et al., 2016; Kourelis et al., 2021). These include the subclades TIR-NLR with the Toll/interleukin-1 receptor (TIR) domain, CC-NLR with the Rx-type coiled-coil (CC) domain, CC_R_-NLR with the RPW8-type CC (CC_R_) domain (Tamborski and Krasileva, 2020) and the more recently defined CC_G10_-NLR with a distinct type of CC (CC_G10_) (Lee et al., 2020). Up to 10% of NLRs carry unconventional “integrated” domains in addition to the canonical tripartite domain architecture. Integrated domains are thought to generally function as decoys to bait pathogen effectors and enable pathogen detection (Cesari et al., 2014; Sarris et al., 2016; Wu et al., 2015; Kourelis and van der Hoorn, 2018). They include dozens of different modules indicating that novel domain acquisitions have repeatedly taken place throughout the evolution of plant NLRs (Sarris et al., 2016; Kroj et al., 2016). To date, over 400 NLRs from 31 genera in 11 orders of flowering plants have been experimentally validated as reported in the RefPlantNLR reference dataset (Kourelis et al., 2021). Several of these NLRs are coded by *R* genes that function against economically important pathogens and contribute to sustainable agriculture (Dangl et al., 2013).

In recent years, the research community has gained a better understanding of the structure/function relationships of plant NLRs and the immune receptor circuitry they form (Wu et al., 2018; Adachi et al., 2019a; Burdett et al., 2019; Jubic et al., 2019; Bayless and Nishimura, 2020; Feehan et al., 2020; Mermigka et al., 2020; Wang and Chai, 2020; Xiong et al., 2020; Zhou and Zhang, 2020). Some NLRs, such as ZAR1, form a single functional unit that carries both pathogen sensing and immune signalling activities in a single protein (termed ‘singleton NLR’ per Adachi et al., 2019a). Other NLRs function together in pairs or more complex networks, where connected NLRs have functionally specialized into sensor NLRs dedicated to pathogen detection or helper NLRs that are required for sensor NLRs to initiate immune signalling (Feehan et al., 2020). Paired and networked NLRs are thought to have evolved from multifunctional ancestral receptors through asymmetrical evolution (Adachi et al., 2019a, 2019b). As a result of their direct coevolution with pathogens, NLR sensors tend to diversify faster than helpers and can be dramatically expanded in some plant taxa (Wu et al., 2017; Stam et al., 2019). For instance, sensor NLRs often exhibit non-canonical biochemical features, such as degenerated functional motifs and unconventional domain integrations (Adachi et al., 2019b; Seong et al., 2020).

The elucidation of plant NLR structures by cryo-electron microscopy has significantly advanced our understanding of the biochemical events associated with the activation of these immune receptors (Wang et al., 2019a; 2019b; Ma et al., 2020; Martin et al., 2020). The CC-NLR ZAR1, the TIR-NLRs RPP1 and Roq1 oligomerize upon activation into a multimeric complex known as the resistosome. In the case of ZAR1, recognition of bacterial effectors occurs through its partner receptor-like cytoplasmic kinases (RLCKs), which occur in a genomic cluster of multiple RLCK-type pseudokinases that vary depending on the pathogen effector and host plant (Lewis et al., 2013; Wang et al., 2015; Seto et al., 2017; Schultink et al. 2019; Laflamme et al., 2020). Activation of ZAR1 induces conformational changes in the nucleotide binding domain resulting in ADP release, dATP/ATP binding and pentamerization of the ZAR1–RLCK complex into the resistosome. The ZAR1 resistosome exposes a funnel-shaped structure formed by the N-terminal α1 helices, which translocates into the plasma membrane, and the resistosome itself acts as a Ca^2+^ channel (Wang et al., 2019b; Bi et al., 2021). The ZAR1 N-terminal α1 helix matches the MADA consensus sequence motif that is functionally conserved in ∼20% of CC-NLRs including NLRs from dicot and monocot plant species (Adachi et al., 2019b). This suggests that the biochemical ‘death switch’ mechanism of the ZAR1 resistosome may apply to a significant fraction of CC-NLRs. Interestingly, unlike singleton and helper CC-NLRs, sensor CC-NLRs often carry degenerated MADA α1 helix motifs and/or N-terminal domain integrations, which would preclude their capacity to trigger cell death according to the ZAR1 model (Adachi et al., 2019b; Seong et al., 2020).

Comparative sequence analyses based on a robust evolutionary framework can yield insights into molecular mechanisms and help generate experimentally testable hypotheses. ZAR1 was previously reported to be conserved across multiple dicot plant species but whether it occurs in other angiosperms hasn’t been systematically studied (Baudin et al. 2017; Schultink et al. 2019; Harant et al. 2021). Here, we used a phylogenomic approach to investigate the molecular evolution of ZAR1 across flowering plants (angiosperms). We discovered 120 ZAR1 orthologs in 88 species, including monocot, magnoliid and eudicot species indicating that ZAR1 is an atypically conserved NLR that traces its origin to early angiosperm lineages ∼220 to 150 million years ago (Jurassic period). We took advantage of this large collection of orthologs to identify highly conserved features of ZAR1, revealing regions for effector recognition, intramolecular interactions and cell death activation. We showed that the cell death activity of ZAR1 from distantly related plant species can be dependent of its partner RLCKs, therefore experimentally validating the hypothesis that ZAR1 has evolved to be a partner with RLCKs early in its evolution. Throughout its evolution, ZAR1 also acquired novel features, including the C-terminal integration of a thioredoxin-like domain and duplication into two paralog families ZAR1-SUB and ZAR1-CIN. Members of the ZAR1-SUB paralog family have highly diversified in eudicots and often lack conserved ZAR1 features. We conclude that ZAR1 has experienced relatively limited gene duplication and expansion throughout its deep evolutionary history, but still did give rise to non-canonical NLR proteins with integrated domains and degenerated molecular features.

## RESULTS

### ZAR1 is the most widely conserved CC-NLR across angiosperms

To determine the distribution of ZAR1 across plant species, we applied a computational pipeline based on iterated BLAST searches of plant genome and protein databases (Figure 1A). These comprehensive searches were seeded with previously identified ZAR1 sequences from Arabidopsis, *N. benthamiana*, tomato, sugar beet and cassava (Baudin et al. 2017; Schultink et al. 2019; Harant et al. 2021). We also performed iterated phylogenetic analyses using the NB-ARC domain of the harvested ZAR1-like sequences, and obtained a well-supported clade that includes the previously reported ZAR1 sequences, as well as new clade members from more distantly related plant species, notably *Colacasia esculenta* (taro, Alismatales), *Cinnamomum micranthum* (Syn. *C. kanehirae*, stout camphor, Magnoliidae) and *Aquilegia coerulea* (columbine, Ranunculales) (Supplemental Data Set 1). In total, we identified 120 ZAR1 from 88 angiosperm species that tightly clustered in the ZAR1 phylogenetic clade (Figure 1B, Supplemental Data Set 1). Among the 120 genes, 108 code for canonical CC-NLR proteins with 52.0 to 97.0% similarity to Arabidopsis ZAR1, whereas another 9 carry the three major domains of CC-NLR proteins but have a C-terminal integrated domain (ZAR1-ID, see below). The remaining 3 genes code for two truncated NLRs and a potentially mis-annotated coding sequence due to a gap in the genome sequence. In summary, we propose that the identified clade consists of ZAR1 orthologs from a diversity of angiosperm species. Our analyses of ZAR1-like sequences also revealed two well-supported sister clades of the ZAR1 ortholog clade (Figure 1B). We named these subclades ZAR1-SUB and ZAR1-CIN and we describe them in more details below.

**Figure 1.**
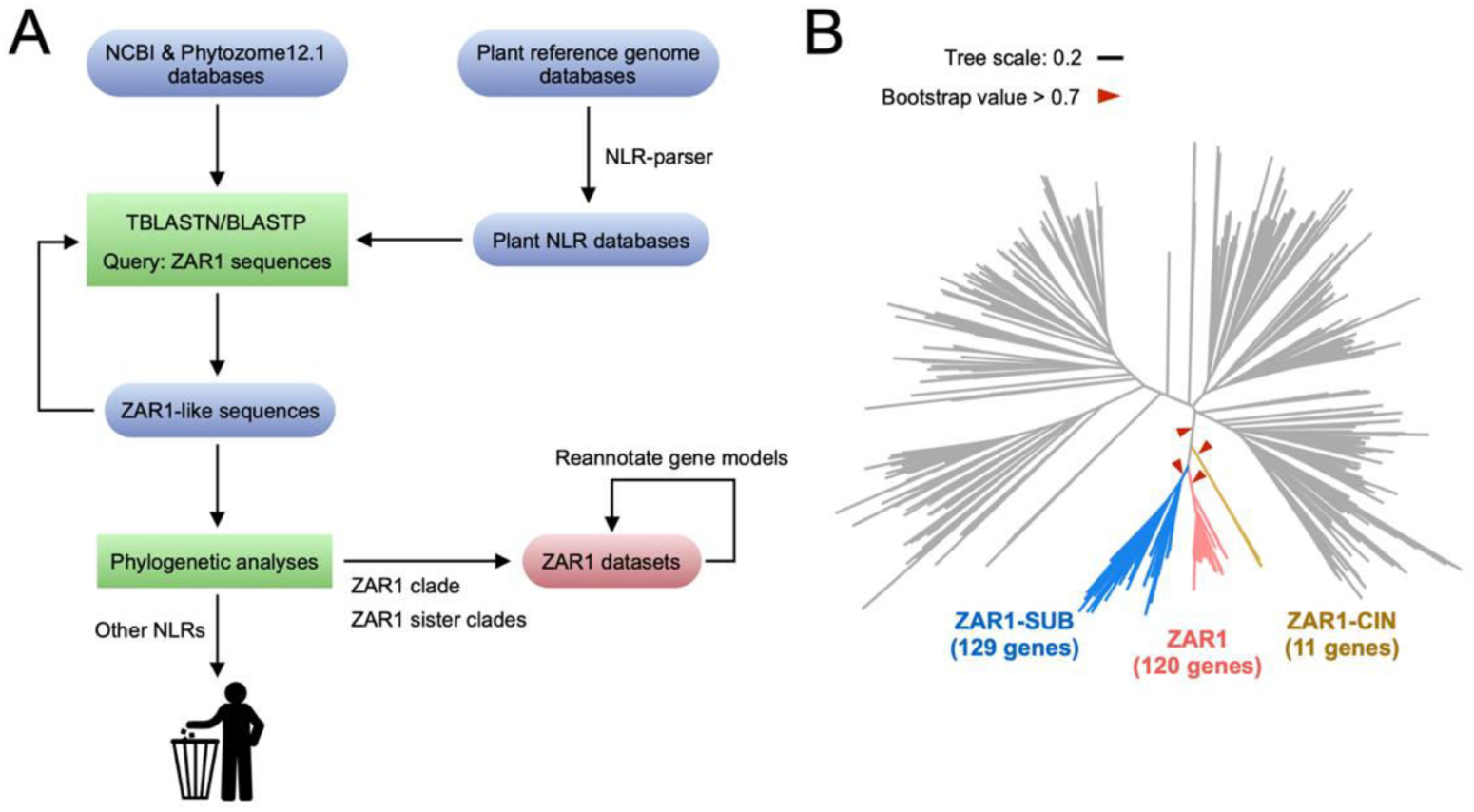
Comparative sequence analyses identify and classify ZAR1 sequences from angiosperms. (**A**) Workflow for computational analyses in searching ZAR1 orthologs. We performed TBLASTN/BLASTP searches and subsequent phylogenetic analyses to identify ZAR1 ortholog genes from plant genome/proteome datasets. (**B**) ZAR1 forms a clade with two closely related sister subclades. The phylogenetic tree was generated in MEGA7 by the neighbour-joining method using NB-ARC domain sequences of ZAR1-like proteins identified from the prior BLAST searches and 1019 NLRs identified from 6 representative plant species, taro, stout camphor, columbine, tomato, sugar beet and Arabidopsis. Each branch is marked with different colours based on the ZAR1 and the sister subclades. Red arrow heads indicate bootstrap support > 0.7 and is shown for the relevant nodes. The scale bar indicates the evolutionary distance in amino acid substitution per site.

We have recently proposed that ZAR1 is the most conserved CC-NLR between rosid and asterid plants (Harant et al. 2021). To further evaluate ZAR1 conservation relative to other CC-NLRs across angiosperms, we used a phylogenetic tree of 1475 NLRs from the monocot taro, the magnoliid stout camphor and 6 eudicot species (columbine, Arabidopsis, cassava, sugar beet, tomato, *N. benthamiana*) to calculate the phylogenetic (patristic) distance between each of the 49 Arabidopsis CC-NLRs and their closest neighbor from each of the other plant species. We found that ZAR1 stands out for having the shortest phylogenetic distance to its orthologs relative to other CC-NLRs in this diverse angiosperm species set (Supplemental Figure 1). A similar analysis where we plotted the phylogenetic distance between each of the 159 *N. benthamiana* CC-NLRs to their closest neighbor from the other species also revealed ZAR1 as displaying the shortest patristic distance across all examined species (Supplemental Figure 2). These analyses revealed that ZAR1 is possibly the most widely conserved CC-NLR in flowering plants (angiosperms).

### Phylogenetic distribution of ZAR1 in angiosperms

Although ZAR1 is distributed across a wide range of angiosperms, we noted particular patterns in its phylogenetic distribution. Supplemental Data Set 1 describes the gene identifiers and other features of ZAR1 orthologs sorted based on the phylogenetic clades reported by Smith and Brown (2018). 68 of the 88 plant species have a single-copy of ZAR1 whereas 20 species have two or more copies (Supplemental Data Set 2). ZAR1 is primarily a eudicot gene but we identified three ZAR1 orthologs outside the eudicots, two in the monocot taro and another one in the magnoliid stout camphor. We failed to detect ZAR1 orthologs in 39 species among the 127 species we examined (Supplemental Data Set 1). Except for taro, ZAR1 is missing in monocot species (17 examined), including in the well-studied *Hordeum vulgare* (barley), *Oryza sativa* (rice), *Triticum aestivum* (wheat) and *Zea mays* (maize). ZAR1 is also missing in all examined species of the eudicot Fabales, Cucurbitales, Apiales and Asterales. However, we found a ZAR1 ortholog in the early diverging eudicot columbine and ZAR1 is widespread in other eudicots, including in 63 rosid, 4 Caryophyllales and 18 asterid species.

### ZAR1 is an ancient Jurassic gene that predates the split between monocots, magnoliids and eudicots

The overall conservation of the 120 ZAR1 orthologs enabled us to perform phylogenetic analyses using the full-length protein sequence and not just the NB-ARC domain as generally done with NLRs (Figure 2, Supplemental Figure 3). These analyses yielded a robust ZAR1 phylogenetic tree with well-supported branches that generally mirrored established phylogenetic relationships between the examined plant species (Smith and Brown, 2018; Chaw et al., 2019). For example, the ZAR1 tree matched a previously published species tree of angiosperms based on 211 single-copy core ortholog genes (Chaw et al., 2019). We conclude that the origin of the ZAR1 gene predates the split between monocots, magnoliids and eudicots and its evolution traced species divergence ever since. We postulate that ZAR1 probably emerged in the Jurassic era ∼220 to 150 million years ago (Mya) based on the species divergence time estimate of Chaw et al. (2019) and consistent with the latest fossil evidence for the emergence of flowering plants (Fu et al., 2018).

**Figure 2.**
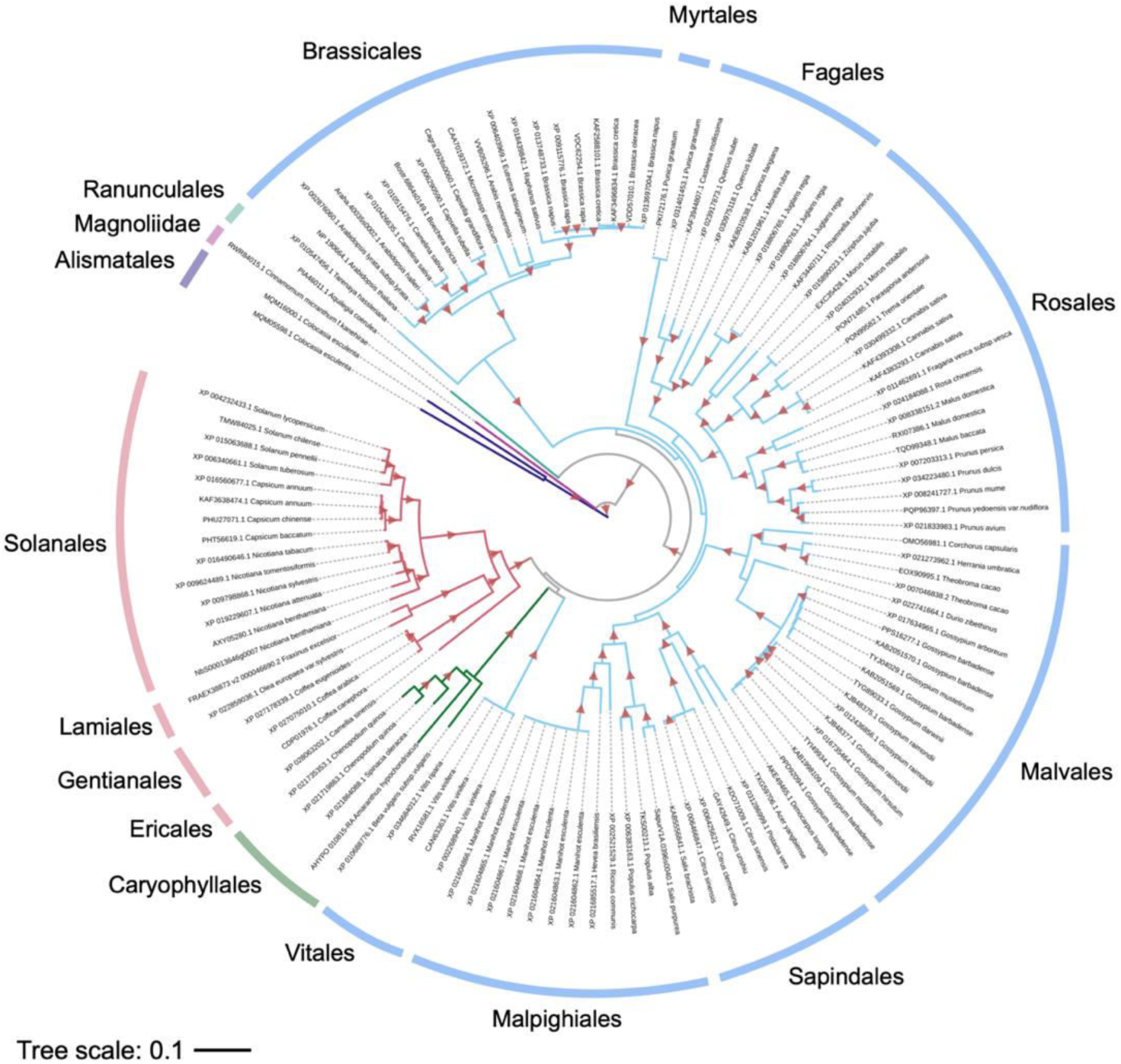
ZAR1 gene is distributed across angiosperms. The phylogenetic tree was generated in MEGA7 by the neighbour-joining method using full length amino acid sequences of 120 ZAR1 orthologs identified in Figure 1. Each branch is marked with different colours based on the plant taxonomy. Red triangles indicate bootstrap support > 0.7. The scale bar indicates the evolutionary distance in amino acid substitution per site.

### ZAR1 is a genetic singleton in a locus that exhibits gene co-linearity across eudicot species

NLR genes are often clustered in loci that are thought to accelerate sequence diversification and evolution (Michelmore and Meyers, 1998; Lee and Chae, 2020). We examined the genetic context of ZAR1 genes using available genome assemblies of taro, stout camphor, columbine, Arabidopsis, cassava, sugar beet, tomato and *N. benthamiana*. The ZAR1 locus is generally devoid of other NLR genes as the closest NLR is found in the Arabidopsis genome 183 kb away from ZAR1 (Supplemental Data Set 3). We conclude that ZAR1 has probably remained a genetic singleton NLR gene throughout its evolutionary history in angiosperms.

Next, we examined the ZAR1 locus for gene co-linearity across the examined species. We noted a limited degree of gene co-linearity between Arabidopsis vs. cassava, cassava vs. tomato, and tomato vs. *N. benthamiana* (Supplemental Figure 4). Flanking conserved genes include the ATPase and protein kinase genes that are present at the ZAR1 locus in both rosid and asterid eudicots. In contrast, we didn’t observe conserved gene blocks at the ZAR1 locus of taro, stout camphor and columbine, indicating that this locus is divergent in these species. Overall, although limited, the observed gene co-linearity in eudicots is consistent with the conclusion that ZAR1 is a genetic singleton with an ancient origin.

### ZAR1 orthologs carry sequence motifs known to be required for Arabidopsis ZAR1 resistosome function

The overall sequence conservation and deep evolutionary origin of ZAR1 orthologs combined with the detailed knowledge of ZAR1 structure and function provide a unique opportunity to explore the evolutionary dynamics of this ancient immune receptor in a manner that cannot be applied to more rapidly evolving NLRs. We used MEME (Multiple EM for Motif Elicitation) (Bailey and Elkan, 1994) to search for conserved sequence patterns among the 117 ZAR1 orthologs (ZAR1 and ZAR1-ID) that encode full-length CC-NLR proteins. This analysis revealed several conserved sequence motifs that span across the ZAR1 orthologs (range of protein lengths: 753-1132 amino acids) (Figure 3A, Supplemental table 1). In Figure 3A, we described the major five sequence motifs or interfaces known to be required for Arabidopsis ZAR1 function that are conserved across ZAR1 orthologs.

**Figure 3.**
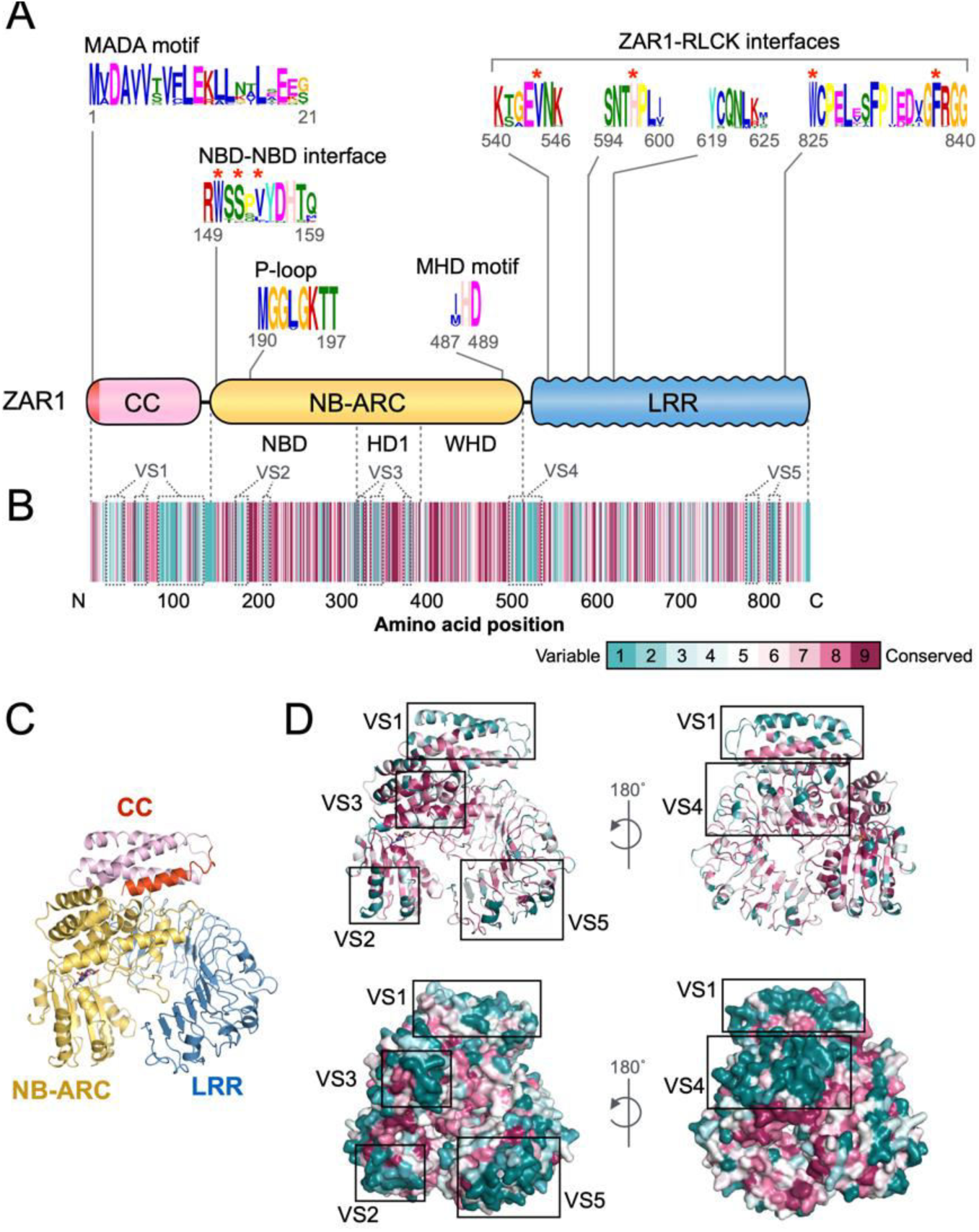
ZAR1 orthologs carry conserved sequence patterns required for Arabidopsis ZAR1 resistosome function. (**A**) Schematic representation of the Arabidopsis ZAR1 protein highlighting the position of conserved sequence patterns across ZAR1 orthologs. Consensus sequence patterns were identified by MEME using 117 ZAR1 ortholog sequences. Raw MEME motifs are listed in Supplemental Table 1. Red asterisks indicate residues functionally validated in Arabidopsis ZAR1 for NBD-NBD and ZAR1-RLCK interfaces. (**B**) Conservation and variation of each amino acid among ZAR1 orthologs across angiosperms. Amino acid alignment of 117 ZAR1 orthologs was used for conservation score calculation via the ConSurf server (https://consurf.tau.ac.il). The conservation scores are mapped onto each amino acid position in Arabidopsis ZAR1 (NP_190664.1). (C, D) Distribution of the ConSurf conservation score on the Arabidopsis ZAR1 structure. The inactive ZAR1 monomer is illustrated in cartoon representation with different colours based on each canonical domain (C) and the conservation score (D). Major five variable surfaces (VS1 to VS5) on the inactive ZAR1 monomer structure are described in grey dot or black boxes in panel B or D, respectively.

Effector recognition by ZAR1 occurs indirectly via binding to RLCKs through the LRR domain. Key residues in the Arabidopsis ZAR1-RLCK interfaces are highly conserved among ZAR1 orthologs and were identified by MEME as conserved sequence patterns (Figure 3A). Valine (V) 544, histidine (H) 597, tryptophan (W) 825 and phenylalanine (F) 839 in the Arabidopsis ZAR1 LRR domain were validated by mutagenesis as important residues for RLCK binding whereas isoleucine (I) 600 was not essential (Wang et al. 2019a; Hu et al. 2020). In the 117 ZAR1 orthologs, V544, H597, W825 and F839 are conserved in 97-100% of the proteins compared to only 63% for I600.

After effector recognition, Arabidopsis ZAR1 undergoes conformational changes from monomeric inactive form to oligomeric active state. This is mediated by ADP release from the NB-ARC domain and subsequent ATP binding, which triggers further structural remodelling in ZAR1 leading to the formation of the activated pentameric resistosome (Wang et al. 2019b). NB-ARC sequences that coordinate binding and hydrolysis of dATP, namely P-loop and MHD motifs, are highly conserved across ZAR1 orthologs (Figure 3A). Histidine (H) 488 and lysine (K) 195, located in the ADP/ATP binding pocket (Wang et al. 2019a; Wang et al. 2019b), are invariant in all 117 orthologs. In addition, three NB-ARC residues, W150, S152 and V154, known to form the NBD-NBD oligomerisation interface for resistosome formation (Wang et al. 2019b; Hu et al. 2020), are present in 82-97% of the ZAR1 orthologs and were also part of a MEME motif (Figure 3A).

The N-terminal CC domain of Arabidopsis ZAR1 mediates cell death signalling thorough the N-terminal α1 helix/MADA motif, that becomes exposed in activated ZAR1 resistosome to form a funnel like structure (Baudin et al., 2017; 2019; Wang et al. 2019b; Adachi et al., 2019b). We detected an N-terminal MEME motif that matches the α1 helix/MADA motif (Figure 3A). We also used the HMMER software (Eddy, 1998) to query the ZAR1 orthologs with a previously reported MADA motif-Hidden Markov Model (HMM) (Adachi et al., 2019b). This HMMER search detected a MADA-like sequence at the N-terminus of all 117 ZAR1 orthologs (Supplemental Data Set 1).

Taken together, based on the conserved motifs depicted in Figure 3A, we propose that angiosperm ZAR1 orthologs share the main functional features of Arabidopsis ZAR1: 1) effector recognition via RLCK binding, 2) remodelling of intramolecular interactions via ADP/ATP switch, 3) oligomerisation via the NBD-NBD interface and 4) α1 helix/MADA motif-mediated activation of hypersensitive cell death.

### ZAR1 resistosome displays conserved surfaces on RLCK binding sites and the inner glutamate ring

To identify additional conserved and variable features in ZAR1 orthologs, we used ConSurf (Ashkenazy et al., 2016) to calculate a conservation score for each amino acid and generate a diversity barcode for ZAR1 orthologs (Figure 3B). The overall pattern is that the 117 ZAR1 orthologs are fairly conserved. Nonetheless, the CC domain (except for the N-terminal MADA motif and a few conserved stretches), the junction between the NB-ARC and LRR domains and the very C-terminus were distinctly more variable than the rest of the protein (Figure 3B).

We also used the cryo-EM structures of Arabidopsis ZAR1 to determine how the ConSurf score map onto the 3D structures (Figure 3C, D and Figure 4). First, we found five major variable surfaces (VS1 to VS5) on the inactive ZAR1 monomer structure (Figure 3C, D), as depicted in the ZAR1 diversity barcode (Figure 3B). VS1 comprises α2/α4 helices and a loop between α3 and α4 helices of the CC domain. VS2 and VS3 corresponds to α1/α2 helices of NBD and a loop between α2 and α3 helices of HD1, respectively. VS4 comprises a loop between WHD and LRR and first three helices of the LRR domain. VS5 is mainly derived from the last three helices of the LRR domain and the loops between these helices (Figure 3B, D).

**Figure 4.**
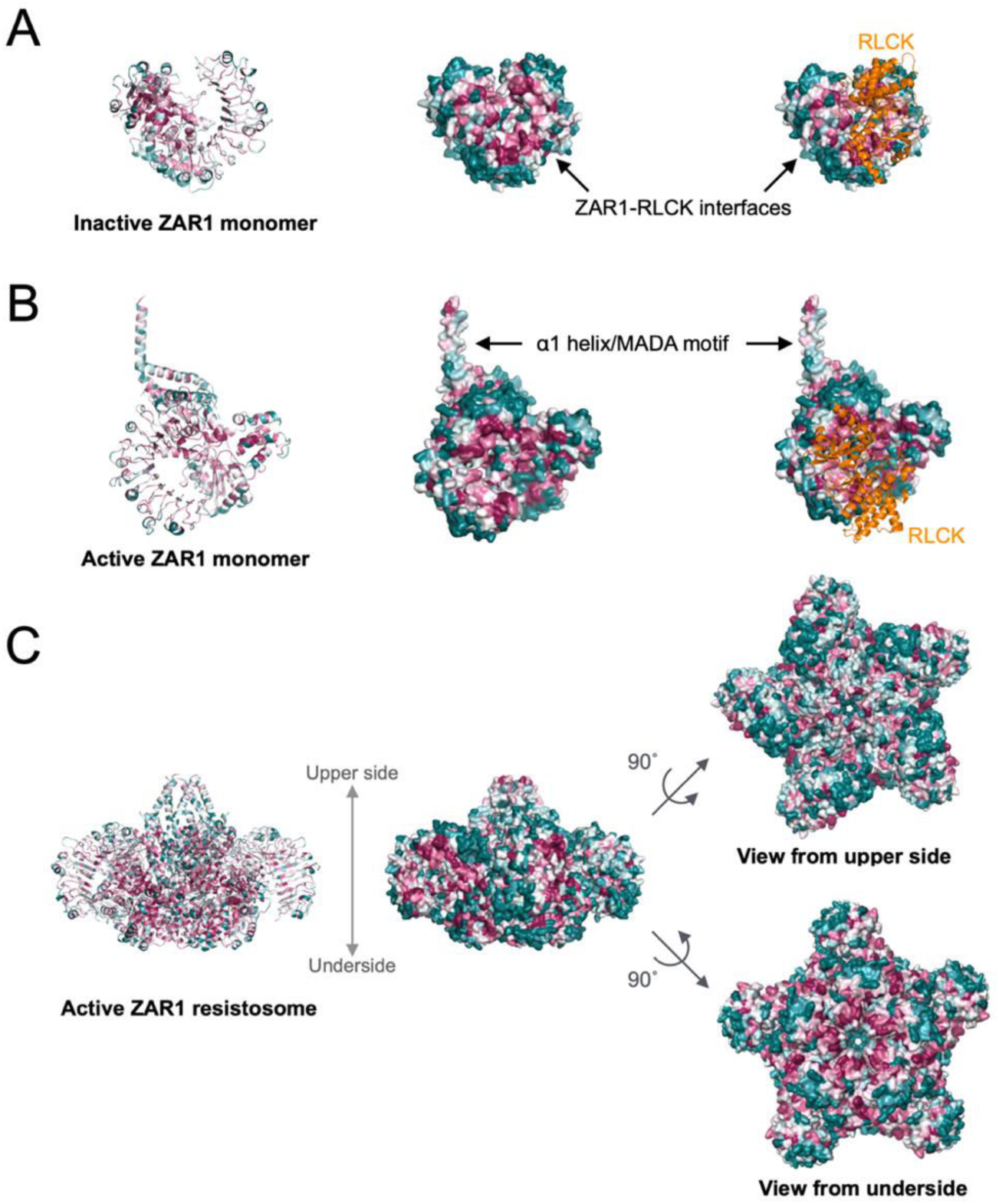
ZAR1 orthologs across angiosperms display multiple conserved surfaces on the resistosome structure. Distribution of the ConSurf conservation score was visualized on the inactive monomer (**A**), active monomer (**B**) and resistosome (**C**) structures of Arabidopsis ZAR1. Each structure and cartoon representation are illustrated with different colours based on the conservation score shown in Figure 3.

Next, we examined highly conserved surfaces on inactive and active ZAR1 structures (Figure 4A, B). Consistent with the MEME analyses, we confirmed that highly conserved surfaces match to the RLCK binding interfaces (Figure 4A, B). We also confirmed that the N-terminal α1 helix/MADA motif is conserved on the resistosome surfaces, although the first four N-terminal amino acids are missing from the N terminus of the active ZAR1 cryo-EM structures (Figure 4B, C). We also noted sequence conservation at the glutamate rings (comprised of E11, E18, E130 and E134) inside the Arabidopsis ZAR1 resistosome (Supplemental Figure 5). Glutamic acid (E) 11 is conserved in 94% of ZAR1 orthologs, whereas only 3-18% retain E18, E130 and E134 in the same positions as Arabidopsis ZAR1. Interestingly, mutation of E11 to alanine (A) impaired Arabidopsis ZAR1-mediated cell death, but the E18A, E130A and E134A mutants were capable of inducing cell death (Bai et al., 2021). Furthermore, the E11A mutation impaired Ca^2+^ channel activity of the ZAR1 resistosome *in vitro* and *in vivo* (Bai et al., 2021). Therefore, our motif and structure analyses suggest that RLCK-mediated effector recognition and E11-dependent Ca^2+^ influx are key functional features conserved across the great majority of ZAR1 orthologs.

### ZAR1 interaction sites are conserved in ZED1-related kinase (ZRK) family proteins across distantly related plant species

We endeavored to experimentally test the hypothesis that ZAR1 ortholog proteins across angiosperm species require RLCKs to activate their molecular switch. First, we searched for RLCK XII-2 subfamily genes in the distantly related plant species, taro, stout camphor and columbine. The BLAST searches of protein databases were seeded with previously identified RLCK ZED1-related kinase (ZRK) sequences from Arabidopsis and *N. benthamiana* (Lewis et al., 2013; Schultink et al. 2019). We also performed iterated phylogenetic analyses using the kinase domain of the harvested ZRK-like sequences and obtained a well-supported clade that includes previously reported ZRK from Arabidopsis (ZRK1∼7, 10∼15) and *N. benthamiana* (JIM2) as well as new clade members from taro, stout camphor and columbine (Figure 5A). In total, we identified 21 ZRK genes in these species, which include 1 ZRK gene (CeZRK1) from taro, 15 ZRK genes (CmZRK1∼15) from stout camphor and 5 ZRK genes (AcZRK1∼5) from columbine (Figure 5A, Supplemental Data Set 4).

**Figure 5.**
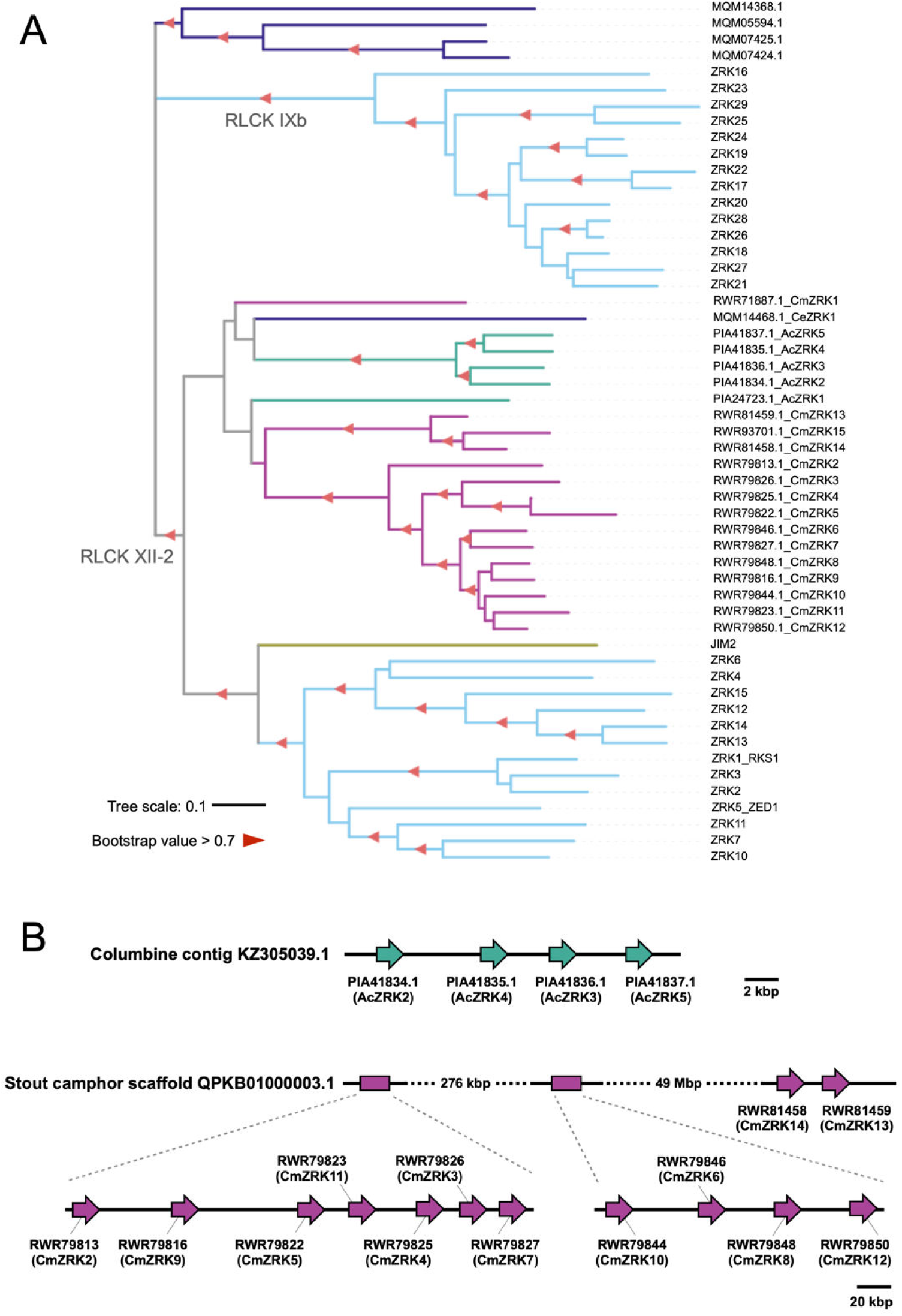
ZRK gene clusters occur in *Aquilegia coerulea* and *Cinnamomum micranthum*. (**A**) The phylogenetic tree was generated in MEGA7 by the neighbour-joining method using full length amino acid sequences of 39 ZRK proteins. Each branch is marked with different colours based on the plant taxonomy. Red triangles indicate bootstrap support > 0.7. The scale bar indicates the evolutionary distance in amino acid substitution per site. (**B**) Schematic representation of the ZRK gene clusters on an A. coerulea (columbine) contig and a *C. micranthum* (Stout camphor) scaffold.

Remarkably, similar to Arabidopsis ZRKs (Lewis et al., 2013), a number of the identified ZRKs are located in genomic clusters. 13 ZRK genes in stout camphor and 4 ZRK genes in columbine form gene clusters on scaffold QPKB01000003.1 and contig KZ305039.1, respectively (Figure 5B). All of the identified ZRK genes are located on different scaffold or contig with ZAR1 gene in taro, stout camphor and columbine, whereas Arabidopsis ZAR1 and the 9 ZRK genes occur on the same chromosome (Supplemental Data Set 4).

The 21 ZRK genes code for proteins of 277-452 amino acids, similar to Arabidopsis and *N. benthamiana* ZRKs, which code for 269-396 amino acid proteins (Supplemental Data Set 5). ZRK family proteins from taro, stout camphor and columbine show 20.8 to 42.2% similarity to Arabidopsis ZRK1 (Supplemental Data Set 6). Although the sequence similarity is low across the ZRK proteins in angiosperms, ZAR1 interaction sites are highly conserved in the ZRKs (Supplemental Figure 6) (Wang et al., 2019a; Hu et al., 2020). Notably, functionally validated residues for ZAR1-RLCK interactions [glycine 27 (G27) and leucine 31 (L31) in Arabidopsis ZRK1; G29 in Arabidopsis ZRK5] are conserved in 81 to 86% of the 21 ZRKs.

Moreover, 90% of the 21 ZRKs have a hydrophobic V or I residue at the same position to V35 in Arabidopsis ZRK1 (corresponding to I24 in Arabidopsis ZRK5). This sequence conservation supports our hypothesis that ZRK family proteins function together with ZAR1 across distantly related plant species.

### Stout camphor and columbine ZED1-related kinase (ZRK) proteins positively regulate ZAR1 autoactive cell death

*N. benthamiana* ZAR1 (NbZAR1) requires its partner RLCK JIM2 to trigger autoimmune cell death in the *N. benthamiana* expression system (Harant et al. 2021). Therefore, we hypothesized that the ZRK family genes are required for ZAR1 to trigger cell death response in angiosperms. To test this hypothesis, we cloned wild-type ZAR1 and ZRK genes from taro, stout camphor and columbine, and also generated autoactive ZAR1 mutants by following the mutation strategy we previously used for the NbZAR1 MHD motif mutant (NbZAR1^D481V^) (Harant et al. 2021). We found that MHD mutants of taro ZAR1 (CeZAR1^D487V^) and columbine ZAR1 (AcZAR1^D489V^), but not stout camphor ZAR1 (CmZAR1^D488V^), induce autoacitive cell death in *N. benthamiana* leaves (Figure 6). Whereas CeZRK1 expression did not alter cell death activity of CeZAR1^D487V^, AcZRK1, AcZRK3, AcZRK4 and AcZRK5, but not AcZRK2, enhanced the cell death response by AcZAR1^D489V^ (Figure 6A to D). Although CmZAR1^D488V^ itself did not trigger visible cell death, coexpression of CmZAR1^D488V^ together with CmZRK2, CmZRK6, CmZRK8, CmZRK9, CmZRK10, CmZRK11 or CmZRK13 caused macroscopic cell death in *N. benthamiana* leaves (Figure 6E, F). These results suggest that ZRKs positively regulate cell death activity of ZAR1 in stout camphor and columbine.

**Figure 6.**
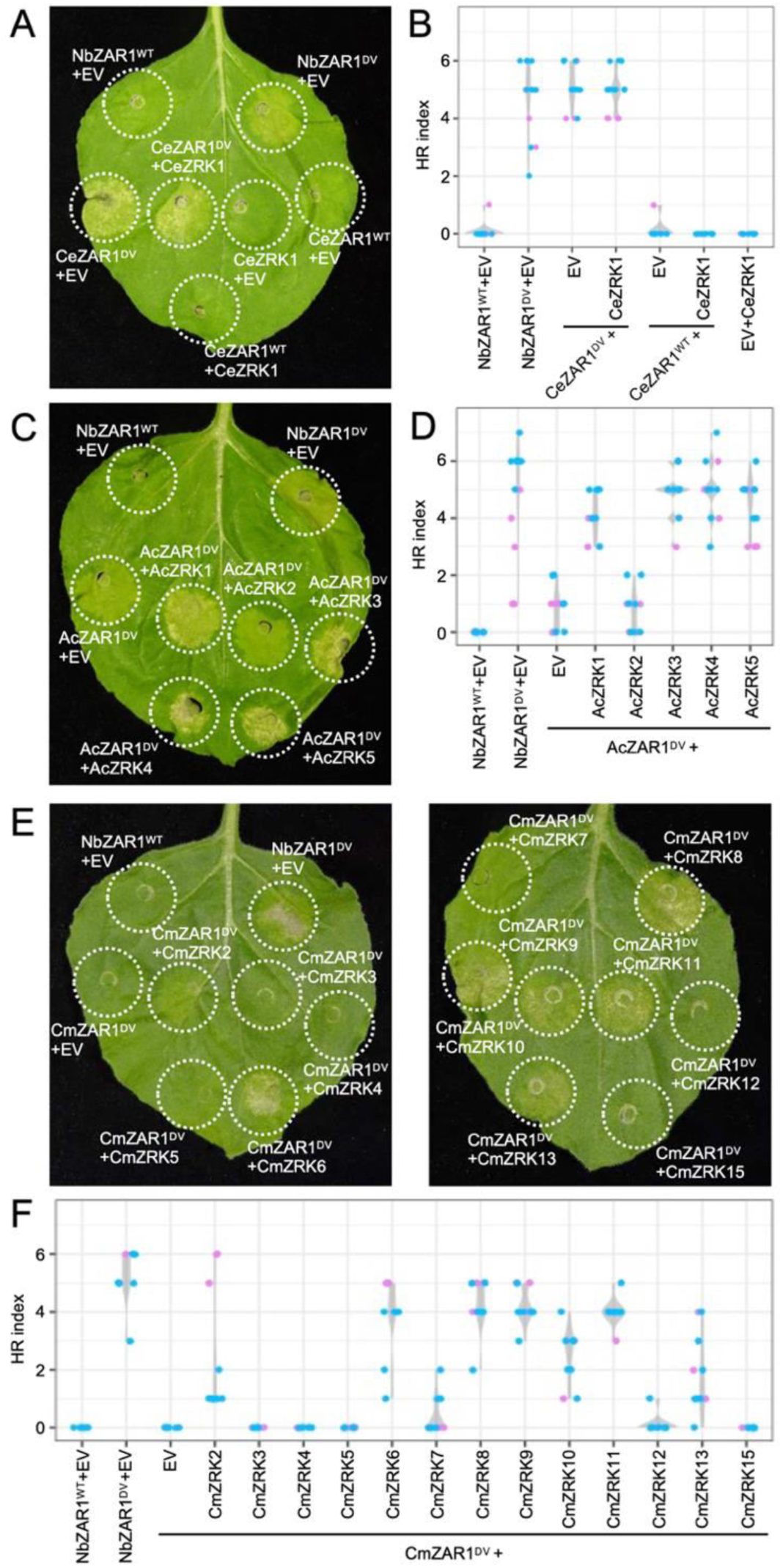
ZRK family proteins positively regulate *Aquilegia coerulea* AcZAR1 and *Cinnamomum micranthum* CmZAR1 autoimmune cell death in Nicotiana benthamiana. (**A, C, E**) Cell death observed in N. benthamiana after expression of ZAR1 mutants with or without wild-type ZRKs. N. benthamiana leaf panels expressing wild-type NbZAR1 (NbZAR1^WT^), NbZAR1^D481V^ (ZAR1^D481V^), wild-type Colacasia esculenta CeZAR1 (CeZAR1^WT^), CeZAR1^D487V^ (CeZAR1^DV^), AcZAR1^D489V^ (AcZAR1^DV^) and CmZAR1^D488V^ (CmZAR1^DV^) with or without wild-type ZRKs, were photographed at 4 to 6 days after agroinfiltration. (**B, D, F**) Violin plots showing cell death intensity scored as an HR index based on three independent experiments.

Our observation that the CeZAR1 autoactive mutant triggered cell death regardless of CeZRK1, raised the possibility that CeZAR1 functions together with the endogenous JIM2 RLCK in *N. benthamiana*. To test this, we used a hairpin-silencing construct of JIM2 (RNAi:JIM2), that mediates silencing of *JIM2* when transiently expressed in *N. benthamiana* leaves (Harant et al. 2021). Silencing of endogenous *JIM2* did not affect the cell death activity of CeZAR1^D487V^, although it suppressed cell death triggered by NbZAR1^D481V^ (Supplemental Figure 7). This result indicates that unlike *N. benthamiana* ZAR1, taro ZAR1 triggers autoactive cell death independently of JIM2.

### Integration of a PLP3a thioredoxin-like domain at the C-termini of cassava and cotton ZAR1

As noted earlier, 9 ZAR1 orthologs carry an integrated domain (ID) at their C-termini (Supplemental Data Set 1). These ZAR1-ID include 2 predicted proteins (XP_021604862.1 and XP_021604864.1) from *Manihot esculenta* (cassava) and 7 predicted proteins (KAB1998109.1, PPD92094.1, KAB2051569.1, TYG89033.1, TYI49934.1, TYJ04029.1, KJB48375.1) from the cotton plant species *Gossypium barbadense*, *Gossypium darwinii*, *Gossypium mustelinum* and *Gossypium raimondii* (Supplemental Data Set 1). The integrations follow an otherwise intact LRR domain and vary in length from 108 to 266 amino acids (Figure 7A). We confirmed that the ZAR1-ID gene models of cassava XP_021604862.1 and XP_021604864.1 are correct based on RNA-seq exon coverage in the NCBI database (database ID: LOC110609538). However, cassava ZAR1-ID XP_021604862.1 and XP_021604864.1 are isoforms encoded by transcripts from a single locus on chromosome LG2 (RefSeq sequence NC_035162.1) of the cassava RefSeq assembly (GCF_001659605.1) which also produces transcripts encoding isoforms lacking the C-terminal ID (XP_021604863.1, XP_021604865.1, XP_021604866.1, XP_021604867.1 and XP_021604868.1). Thus, cassava ZAR1-ID are probably splicing variants from a unique cassava ZAR1 gene locus (Supplemental Figure 8).

**Figure 7.**
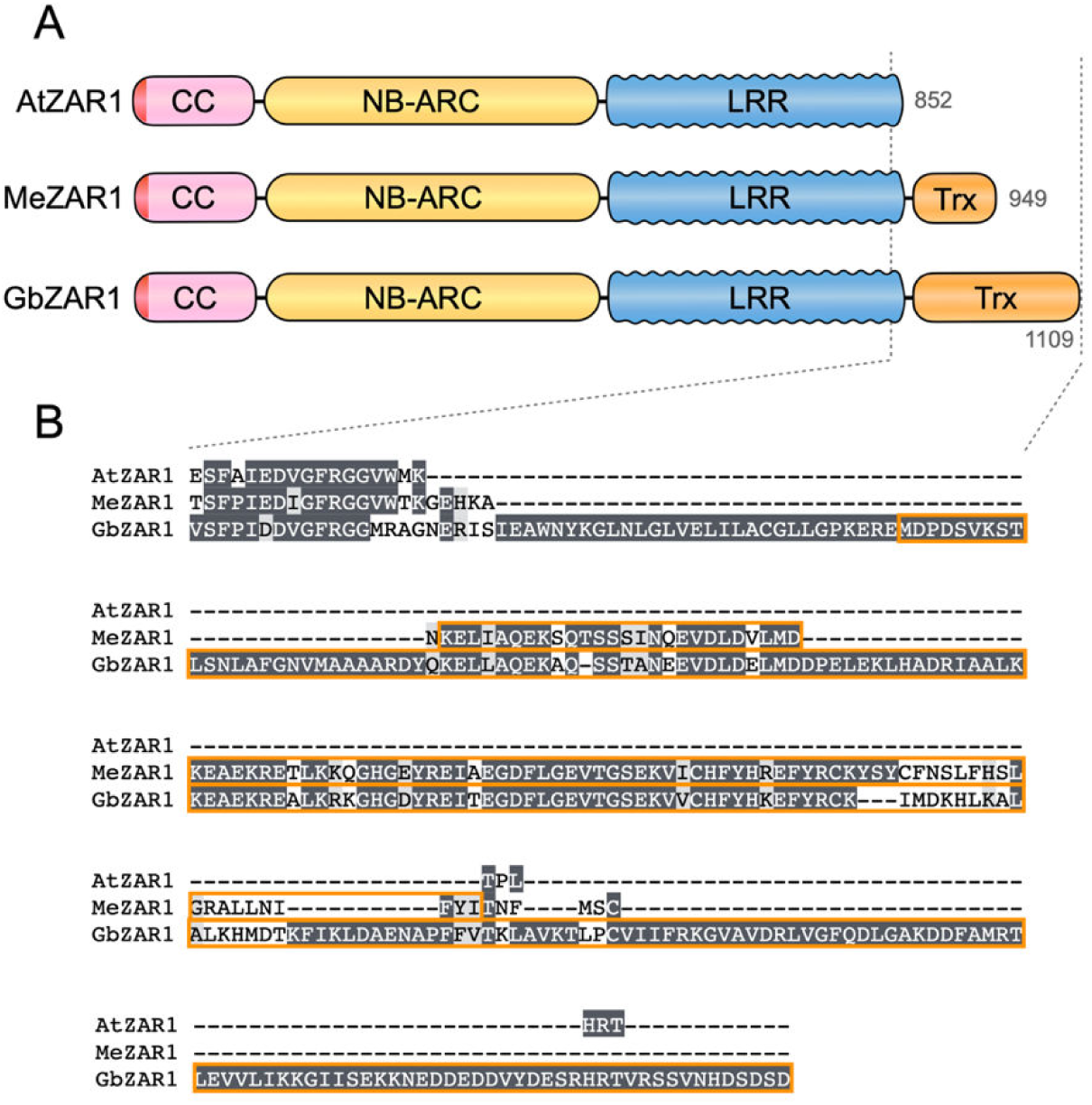
Cassava and cotton ZAR1-ID carry an additional Trx domain at the C terminus. (**A**) Schematic representation of NLR domain architecture with C-terminal Trx domain. (**B**) Description of Trx domain sequences on amino acid sequence alignment. Cassava XP_021604862.1 (MeZAR1) and cotton KAB1998109.1 (GbZAR1) were used for MAFFT version 7 alignment as representative ZAR1-ID. Arabidopsis ZAR1 (AtZAR1) was used as a control of ZAR1 without ID.

To determine the phylogenetic relationship between ZAR1-ID and canonical ZAR1, we mapped the domain architectures of ZAR1 orthologs on the phylogenetic tree shown in Figure 2 (Supplemental Figure 9). Cassava and cotton ZAR1-ID occur in different branches of the ZAR1 rosid clade indicating that they may have evolved as independent integrations although alternative evolutionary scenarios such as a common origin followed by subsequent deletion of the ID or lineage sorting remain possible (Supplemental Figure 9).

We annotated all the C-terminal extensions as thioredoxin-like using InterProScan (Trx, IPR036249; IPR013766; cd02989). The integrated Trx domain sequences share sequence similarity to each other (Figure 7B). They are also similar to Arabidopsis AT3G50960 (phosphoducin-like PLP3a; 34.8-90% similarity to integrated Trx domains), which is located immediately downstream of ZAR1 in a tail-to-tail configuration in the Arabidopsis genome (Supplemental Figure 10). We also noted additional genetic linkage between ZAR1 and Trx genes in other rosid species, namely field mustard, orange, cacao, grapevine and apple, and in the asterid species coffee (Supplemental Data Set 7). We conclude that ZAR1 is often genetically linked to a PLP3a-like Trx domain gene and that the integrated domain in ZAR1-ID has probably originated from a genetically linked sequence.

### The ZAR1-SUB clade emerged early in eudicot evolution from a single ZAR1 duplication event

Phylogenetic analyses revealed ZAR1-SUB as a sister clade of the ZAR1 ortholog clade (Figure 1B, Figure 8). ZAR1-SUB clade comprises 129 genes from a total of 55 plant species (Supplemental Data Set 8). 21 of the 55 plant species carry a single-copy of ZAR1-SUB whereas 34 species have two or more copies (Supplemental Data Set 2). Of the 129 genes, 122 code for canonical CC-NLR proteins (692-1038 amino acid length) with shared sequence similarities ranging from 36.5 to 99.9% (Supplemental Data Set 8).

**Figure 8.**
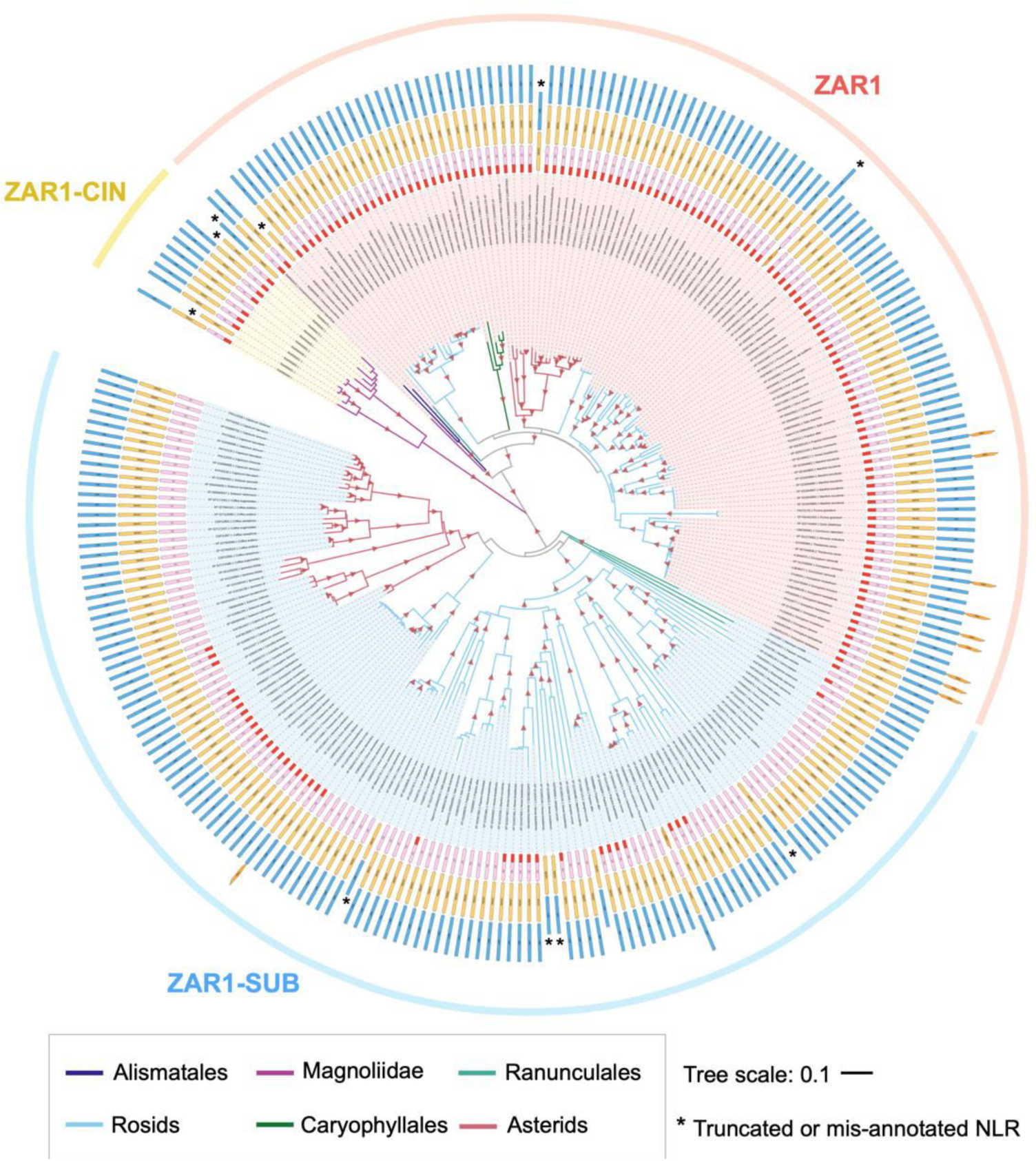
ZAR1-SUB has emerged early in eudicots and diverged at MADA motif sequence. The phylogenetic tree was generated in MEGA7 by the neighbour-joining method using full length amino acid sequences of 120 ZAR1, 129 ZAR1-SUB and 11 ZAR1-CIN identified in Figure 1. Each branch is marked with different colours based on the plant taxonomy. Red triangles indicate bootstrap support > 0.7. The scale bar indicates the evolutionary distance in amino acid substitution per site. NLR domain architectures are illustrated outside of the leaf labels: MADA is red, CC is pink, NB-ARC is yellow, LRR is blue and other domain is orange. Black asterisks on domain schemes describe truncated NLRs or potentially mis-annotated NLR.

Unlike ZAR1, ZAR1-SUB NLRs are restricted to eudicots (Supplemental Figure 11, Supplemental Data Set 8). Three out of 129 genes are from the early diverging eudicot clade Ranunculales species, namely columbine, *Macleaya cordata* (plume poppy) and *Papaver somniferum* (opium poppy). The remaining ZAR1-SUB are spread across rosid and asterid species. We found that 11 species have ZAR1-SUB genes but lack a ZAR1 ortholog (Supplemental Data Set 2). These 11 species include two of the early diverging eudicots plume poppy and opium poppy, and the Brassicales *Carica papaya* (papaya). Interestingly, papaya is the only Brassicales species carrying a ZAR1-SUB gene, whereas the 16 other Brassicales species have ZAR1 but lack ZAR1-SUB genes (Supplemental Data Set 2). In total, we didn’t detect ZAR1-SUB genes in 44 species that have ZAR1 orthologs, and these 44 species include the monocot taro, the magnoliid stout camphor and 42 eudicots, such as Arabidopsis, sugar beet and *N. benthamiana* (Supplemental Data Set 2).

In summary, given the taxonomic distribution of the ZAR1-SUB clade genes, we propose that ZAR1-SUB has emerged from a single duplication event of ZAR1 prior to the split between Ranunculales and other eudicot lineages about ∼120-130 Mya based on the species divergence time estimate of Chaw et al. (2019).

### ZAR1-SUB paralogs have significantly diverged from ZAR1

We investigated the sequence patterns of ZAR1-SUB proteins and compared them to the sequence features of canonical ZAR1 proteins that we identified earlier (Figure 3A). MEME analyses revealed several conserved sequence motifs (Supplemental Table 2). Especially, the MEME motifs in the ZAR1-SUB NB-ARC domain were similar to ZAR1 ortholog motifs (Supplemental Table 3). These include P-loop and MHD motifs, which are broadly conserved in NB-ARC of 97% and 100% of the ZAR1-SUB NLRs, respectively (Figure 9A). MEME also revealed sequence motifs in the ZAR1-SUB LRR domain that partially overlaps in position with the conserved ZAR1-RLCK interfaces (Figure 9A, Supplemental Figure 12). However, the ZAR1-SUB MEME motifs in the LRR domain were variable at the ZAR1-RLCK interface positions compared to ZAR1, and the motif sequences were markedly different between ZAR1-SUB and ZAR1 proteins (Figures 3A, 9A).

**Figure 9.**
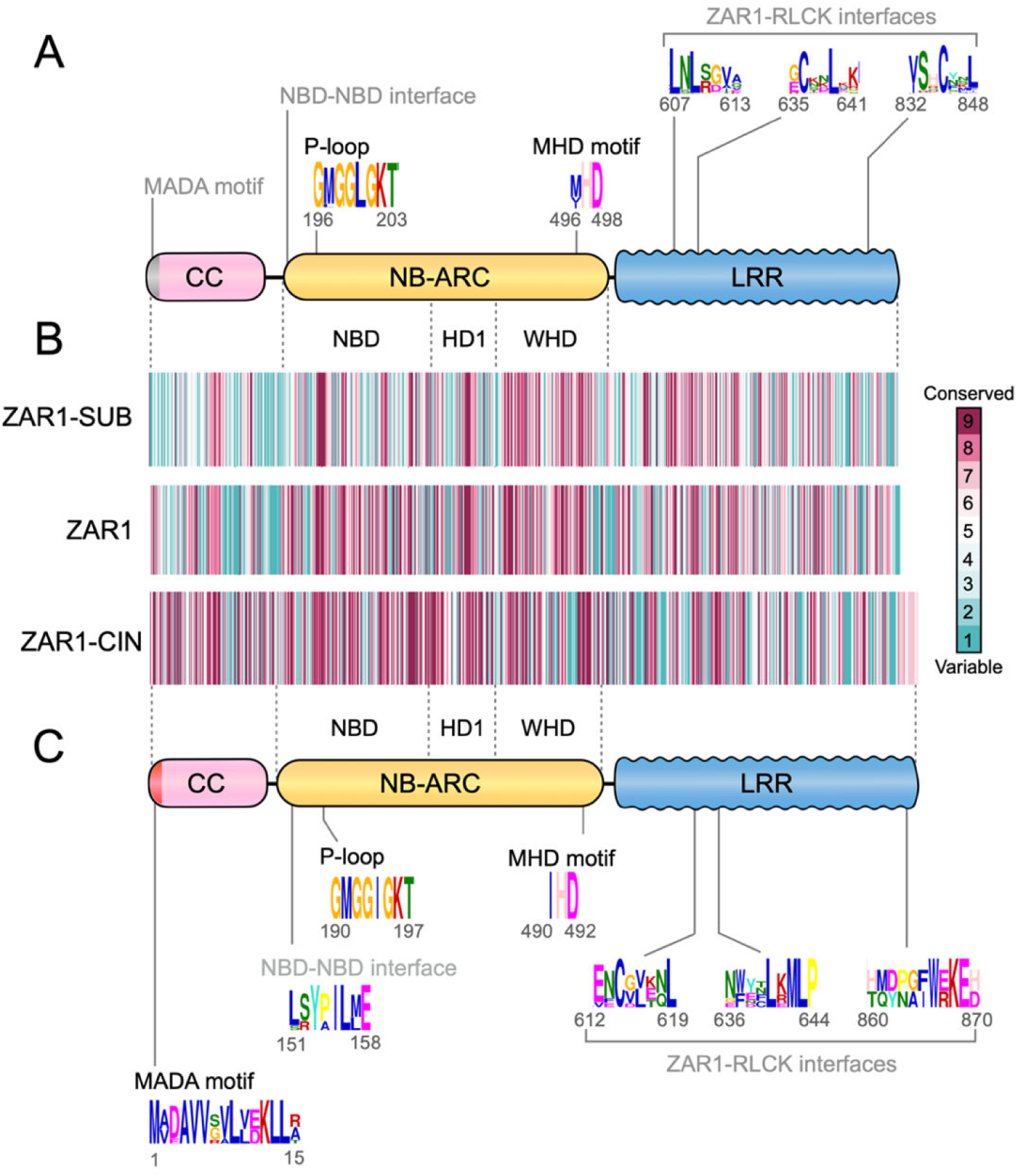
Conserved sequence distributions in ZAR1-SUB and ZAR1-CIN. (**A**) Schematic representation of the ZAR1-SUB protein highlighting the position of the representative conserved sequence patterns across ZAR1-SUB. Representative consensus sequence patterns identified by MEME are described on the scheme. Raw MEME motifs are listed in Supplemental Tables 2 and 3. (**B**) Conservation and variation of each amino acid among ZAR1-SUB and ZAR1-CIN. Amino acid alignment of 129 ZAR1-SUB or 8 ZAR1-CIN was used for conservation score calculation via the ConSurf server (https://consurf.tau.ac.il). The conservation scores are mapped onto each amino acid position in queries XP_004243429.1 (ZAR1-SUB) and RWR85656.1 (ZAR1-CIN), respectively. (**C**) Schematic representation of the ZAR1-CIN protein highlighting the position of the representative conserved sequence patterns across 8 ZAR1-CIN. Raw MEME motifs are listed in Supplemental Tables 4 and 5.

Remarkably, unlike ZAR1 orthologs, MEME did not predict conserved sequence pattern from a region corresponding to the MADA motif, indicating that these sequences have diverged across ZAR1-SUB proteins (Figure 9A). We confirmed the low frequency of MADA motifs in ZAR1-SUB proteins using HMMER searches with only ∼30% (38 out of 129) of the tested proteins having a MADA-like sequence (Supplemental Data Set 8, Figure 8). Moreover, conserved sequence patterns were not predicted for the NBD-NBD interface and the conserved underside surface of the ZAR1 resistosome (Figure 9A, Supplemental Figure 12). This indicates that the NB-ARC domain of ZAR1-SUB proteins is highly diversified in contrast to the relatively conserved equivalent region of ZAR1 proteins.

We generated a diversity barcode for ZAR1-SUB proteins using the ConSurf as we did earlier with ZAR1 orthologs (Figure 9B). This revealed that there are several conserved sequence blocks in each of the CC, NB-ARC and LRR domains, such as the regions corresponding to P-loop, MHD motif and the equivalent of the ZAR1-RLCK interfaces. Nonetheless, ZAR1-SUB proteins are overall more diverse than ZAR1 orthologs especially in the CC domain, including the N-terminal MADA motif, and the NBD/HD1 regions of the NB-ARC domain where the NBD-NBD interface is located.

Next, we mapped the ConSurf conservation scores onto a homology model of a representative ZAR1-SUB protein (XP_004243429.1 from tomato) built based on the Arabidopsis ZAR1 cryo-EM structures (Supplemental Figure 13). As highlighted in Supplemental Figure 13B and C, conserved residues, such as MHD motif region in the WHD, are located inside of the monomer and resistosome structures. Interestingly, although the prior MEME prediction analyses revealed conserved motifs in positions matching the ZAR1-RLCK interfaces in the LRR domain, the ZAR1-SUB structure homology models displayed variable surfaces in this region (Supplemental Figure 13A). This indicates that the variable residues within these sequence motifs are predicted to be on the outer surfaces of the LRR domain and may reflect interaction with different ligands.

Taken together, these results suggest that unlike ZAR1 orthologs, the ZAR1-SUB paralogs have divergent molecular patterns for regions known to be involved in effector recognition, resistosome formation and activation of hypersensitive cell death.

### Eleven tandemly duplicated ZAR1-CIN genes occur in a 500 kb cluster in the *Cinnamomum micranthum* (stout camphor) genome

The ZAR1-CIN clade, identified by phylogenetic analyses as a sister clade to ZAR1 and ZAR1-SUB, consists of 11 genes from the magnoliid species stout camphor (Figure 1B, Figure 8, Supplemental Data Set 9). 8 of the 11 ZAR1-CIN genes code for canonical CC-NLR proteins with 63.8 to 98.9% sequence similarities to each other, whereas the remaining 3 genes code for truncated NLR proteins. Interestingly, all ZAR1-CIN genes occur in a ∼500 kb cluster on scaffold QPKB01000005.1 of the stout camphor genome assembly (GenBank assembly accession GCA_003546025.1) (Supplemental Figure 14A, B). This scaffold also contains the stout camphor ZAR1 ortholog (CmZAR1, RWR84015), which is located 48 Mb from the ZAR1-CIN cluster (Supplemental Figure 14 A, B). Based on the observed phylogeny and gene clustering, we suggest that the ZAR1-CIN cluster emerged from segmental duplication and expansion of the ancestral ZAR1 gene after stout camphor split from the other examined ZAR1 containing species.

We examined the expression of the eleven CmZAR1 and ZAR1-CIN genes in seven tissues of **C. micranthum** based on the data of Chaw et al. (Chaw et al., 2019). The CmZAR1 gene is relatively highly expressed in seven different tissues of the stout camphor tree (Supplemental Figure 14C). In contrast, only five of the eleven ZAR1-CIN genes displayed detectable expression levels. Of these, two ZAR1-CIN genes (RWR85656 and RWR85657) had different expression patterns across the tissues. Whereas RWR85657 had the highest expression level in flowers, RWR85656 displayed the highest expression levels in stem and old leaf tissues (Supplemental Figure 14C). The implications of these observations remain unclear but may reflect different degrees of tissue specialization of the ZAR1-CIN genes.

### Tandemly duplicated ZAR1-CIN display variable ligand binding interfaces on the LRR domain

We performed MEME and ConSurf analyses of the 8 intact ZAR1-CIN proteins as described above for ZAR1 and ZAR1-SUB. The ConSurf barcode revealed that although ZAR1-CIN proteins are overall conserved, their WHD region and LRR domain include some clearly variable blocks (Figure 9B). MEME analyses of ZAR1-CIN sequences revealed that like ZAR1 orthologs, the MADA, P-loop and MHD motifs match highly conserved blocks of the ZAR1-CIN ConSurf barcode (Figure 9B, C, Supplemental Tables 4 and 5). Consistently, 87.5% (7 out of 8) of the ZAR1-CIN proteins were predicted to have a MADA-type N-terminal sequence based on MADA-HMM analyses (Supplemental Data Set 9, Figure 8).

MEME picked up additional sequence motifs in ZAR1-CIN proteins that overlap in position with the NBD-NBD and ZAR1-RLCK interfaces (Figure 9C, Supplemental Figure 15). However, the sequence consensus at the NBD-NBD and ZAR1-RLCK interfaces indicated these motifs are more variable among ZAR1-CIN proteins relative to ZAR1 orthologs, and the motif sequences were markedly different from the matching region in ZAR1 (Figures 3A, 9C).

We also mapped the ConSurf conservation scores onto a homology model of a representative ZAR1-CIN protein (RWR85656.1) built based on the Arabidopsis ZAR1 cryo-EM structures (Supplemental Figure 13). This model revealed several conserved surfaces, such as on the α1 helix in the CC domain and the WHD of the NB-ARC domain (Supplemental Figure 13B, C). In contrast, the ZAR1-CIN structure homology models displayed highly varied surfaces especially in the LRR region matching the RLCK binding interfaces of ZAR1 (Supplemental Figure 13A). This sequence diversification on the LRR surface suggests that the ZAR1-CIN paralogs may have different host partner proteins and/or effector recognition specificities compared to ZAR1.

## DISCUSSION

This study of ZAR1 macroevolution originated from phylogenomic analyses we initiated during the UK COVID-19 lockdown of March 2020. We performed iterated comparative sequence similarity searches of plant genomes using the CC-NLR immune receptor ZAR1 as a query, and subsequent phylogenetic evaluation of the recovered ZAR1-like sequences. This revealed that ZAR1 is an ancient gene with 120 orthologs recovered from 88 species including monocot, magnoliid and eudicot plants. ZAR1 is an atypically conserved NLR in these species with the gene phylogeny tracing species phylogeny, and consistent with the view that ZAR1 originated early in angiosperms during the Jurassic geologic period ∼220 to 150 Mya (Figure 10). The ortholog series enabled us to determine that resistosome sequences that are known to be functionally important and have remained highly conserved throughout the long evolutionary history of ZAR1. In addition, we experimentally validated the model that ZAR1 has been a partner with RLCKs for over 150 Mya through functional reconstruction of ZAR1-RLCK pairs from distantly related plant species (Figure 6). The main unexpected feature among ZAR1 orthologs is the acquisition of a C-terminal thioredoxin-like domain in cassava and cotton species (Figure 7). Our phylogenetic analyses also indicated that ZAR1 duplicated twice throughout its evolution (Figure 10). In the eudicots, ZAR1 spawned a large paralog family, ZAR1-SUB, which greatly diversified and often lost the typical sequence features of ZAR1. A second paralog, ZAR1-CIN, is restricted to a tandemly repeated 11-gene cluster in stout camphor. Overall, our findings map patterns of functional conservation, expansion and diversification onto the evolutionary history of ZAR1 and its paralogs.

**Figure 10.**
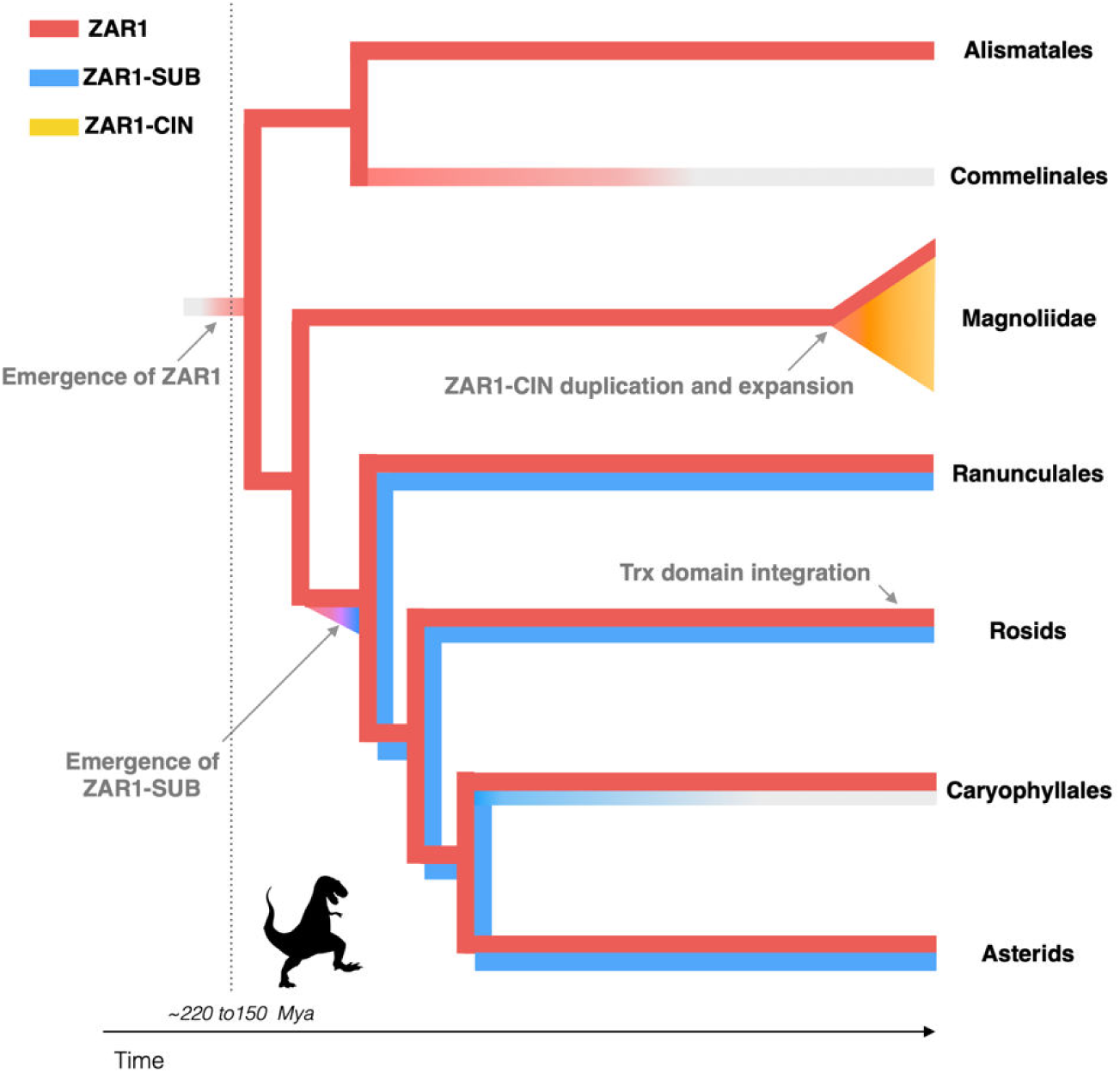
Evolution of ZAR1 and the paralogs in angiosperms. We propose that the ancestral ZAR1 gene has emerged ∼220 to 150 million years ago (Mya) before monocot and eudicot lineages split. ZAR1 gene is widely conserved CC-NLR in angiosperms, but it is likely that ZAR1 has lost in a monocot lineage, Commelinales. A sister clade paralog ZAR1-SUB has emerged early in the eudicot lineages and may have lost in Caryophyllales. Another sister clade paralog ZAR1-CIN has duplicated from ZAR1 gene and expanded in the Magnoliidae *C. micranthum*. Trx domain integration to C terminus of ZAR1 has independently occurred in few rosid lineages.

ZAR1 most likely emerged prior to the split between monocots, Magnoliids and eudicots, which corresponds to ∼220 to 150 Mya based on the dating analyses of Chaw et al. (2019). The origin of the angiosperms remains hotly debated with uncertainties surrounding some of the fossil record coupled with molecular clock analyses that would benefit from additional genome sequences of undersampled taxa (Coiro et al., 2019). Recently, Fu et al. (2018) provided credence to an earlier emergence of angiosperms with the discovery of the fossil flower *Nanjinganthus dendrostyla*, which places the emergence of flowering plants at the Early Jurassic. It is tempting to speculate that ZAR1 emerged among these early flowering plants during the period when dinosaurs dominated planet earth.

NLRs are notorious for their rapid and dynamic evolutionary patterns even at the intraspecific level. In sharp contrast, ZAR1 is an atypical core NLR gene conserved in a wide range of angiosperm species (Figure 2). Nevertheless, Arabidopsis ZAR1 can recognize diverse bacterial pathogen effectors, including five different effector families distributed among nearly half of a collection of ∼500 *Pseudomonas syringae* strains (Laflamme et al., 2020) and an effector AvrAC from *Xanthomonas campestris* (Wang et al., 2015). How did ZAR1 remain conserved throughout its evolutionary history while managing to detect a diversity of effectors? The answer to the riddle lies in the fact that ZAR1 effector recognition occurs via its partner RLCKs. ZRKs of the RLCK XII-2 subfamily rest in complex with inactive ZAR1 proteins and bait effectors by binding them directly or by recruiting other effector-binding RLCKs, such as the family VII PBS1-like protein 2 (PBL2) (Lewis et al., 2013; Wang et al., 2015). These ZAR1-associated RLCKs are highly diversified not only in Arabidopsis (Lewis et al., 2013), but also in stout camphor and columbine, where RLCK XII-2 members occurring in expanded ZRK gene clusters (Figure 5). In the Arabidopsis ZRK cluster, RKS1/ZRK1 is required for recognition of *X. campestris* effector AvrAC (Wang et al., 2015) and ZRK3 and ZRK5/ZED1 are required for recognition of *P. syringae* effectors HopF2a and HopZ1a, respectively (Lewis et al., 2013; Seto et al., 2017). Therefore, as in the model discussed by Schultink et al. (2019), RLCKs appear to have evolved as pathogen ‘sensors’ whereas ZAR1 acts as a conserved signal executor to activate immune response.

The MEME and ConSurf analyses are consistent with the model of ZAR1/RLCK evolution described above. ZAR1 is not just exceptionally conserved across angiosperms but it has also preserved sequence patterns that are key to resistosome-mediated immunity (Figures 3 and 4). Within the LRR domain, ZAR1 orthologs display highly conserved surfaces for RLCK binding (Figure 4). We conclude that ZAR1 has been guarding host kinases throughout its evolution ever since the Jurassic period. These findings strikingly contrast with observations recently made by Prigozhin and Krasileva (2020) on highly variable Arabidopsis NLRs (hvNLRs), which tend to have diverse LRR sequences. For instance, the CC-NLR RPP13 displays variable LRR surfaces across 62 Arabidopsis accessions, presumably because these regions are effector recognition interfaces that are caught in arms race coevolution with the oomycete pathogen *Hyaloperonospora arabidopsidis* (Prigozhin and Krasileva, 2020). The emerging view is that the mode of pathogen detection (direct vs indirect recognition) drives an NLR evolutionary trajectory by accelerating sequence diversification at the effector binding site or by maintaining the binding interface with the partner guardee/decoy proteins (Prigozhin and Krasileva, 2020).

Our functional validation of ZAR1 and ZRKs from distantly related plant species supported the model that ZRKs function together with ZAR1 to trigger immune response *in planta* (Figure 6). 11 of the 19 tested ZRKs were either required or enhanced ZAR1 autoactivity in *N. benthamiana*. The remaining 8 tested ZRKs, CeZRK1, AcZRK2, CmZRK3, CmZRK4, CmZRK5, CmZRK7, CmZRK12 and CmZRK15 did not alter cell death activity of ZAR1. Notably, CmZRK4, CmZRK5 and CmZRK12 have N-terminal truncation or mutations at the ZAR1 interaction sites identified from the Arabidopsis ZAR1-ZRK studies (Supplemental Data Set 6). Therefore, some of the ZRK members may have lost their association with ZAR1 through deletion or mutations. Taro and columbine ZAR1 could trigger autoactive cell death without their partner RLCKs in *N. benthamiana*, whereas stout camphor and *N. benthamiana* ZAR1 proteins require ZRKs to trigger the cell death response (Figure 6; Supplemental Figure 7) (Harant et al., 2021). In the case of taro and columbine, ZRKs may trigger conformational changes of ZAR1 after recognition of cognate pathogen effectors. In this scenario, autoactive ZAR1 could form a resistosome without ZRK proteins, thereby triggering the observed cell death response. In the future, further comparative biochemical studies would further inform our understanding of how ZAR1-ZRK interactions contribute to resistosome formation across angiosperms.

ZAR1 orthologs display a patchy distribution across angiosperms (Supplemental Data Set 1). Given the low number of non-eudicot species with ZAR1 it is challenging to develop a conclusive evolutionary model. Nonetheless, the most parsimonious explanation is that ZAR1 was lost in the monocot Commelinales lineage (Figure 10, Supplemental Data Set 1). ZAR1 is also missing in some eudicot lineages, notably Fabales, Cucurbitales, Apiales and Asterales (Supplemental Data Set 1). Cucurbitaceae (Cucurbitales) species are known to have reduced repertoires of NLR genes possibly due to low levels of gene duplications and frequent deletions (Lin et al., 2013). ZAR1 may have been lost in this and other plant lineages as part of an overall shrinkage of their NLRomes or as consequence of selection against autoimmune phenotypes triggered by NLR mis-regulation (Karasov et al., 2017; Adachi et al., 2019a). In the future, it would be interesting to investigate the repertoires of RLCK subfamilies VII and XII in species that lack ZAR1 orthologs.

We unexpectedly discovered that some ZAR1 orthologs from cassava and cotton species carry a C-terminal thioredoxin-like domain (ZAR1-ID in Figure 7). What is the function of these integrated domains? The occurrence of unconventional domains in NLRs is relatively frequent and ranges from 5 to 10% of all NLRs. In several cases, integrated domains have emerged from pathogen effector targets and became decoys that mediate detection of the effectors (Kourelis and van der Hoorn, 2018). Whether or not the integrated Trx domain of ZAR1-ID functions to bait effectors will need to be investigated. Since ZAR1-ID proteins still carry intact RLCK binding interfaces (Supplemental Data Set 10), they may have evolved dual or multiple recognition specificities via RLCKs and the Trx domain. In addition, all ZAR1-ID proteins have an intact N-terminal MADA motif (Supplemental Figure 9), suggesting that they probably can execute the hypersensitive cell death through their N-terminal CC domains even though they carry a C-terminal domain extension (Adachi et al., 2019b). However, we noted multiple splice variants of the ZAR1-ID gene of cassava, some of which lack the Trx integration (Supplemental Figure 8). It is possible that both ZAR1 and ZAR1-ID isoforms are produced, potentially functioning together as a pair of sensor and helper NLRs.

Our sequence analyses of ZAR1-ID indicate that the integrated Trx domain originates from the PLP3 phosphoducin gene, which is immediately downstream of ZAR1 in the Arabidopsis genome and adjacent to ZAR1 in several other eudicot species (Supplemental Figure 10). Whether or not PLP3 plays a role in ZAR1 function and the degree to which close genetic linkage facilitated domain fusion between these two genes are provocative questions for future studies.

ZAR1 spawned two classes of paralogs through two independent duplication events. The ZAR1-SUB paralog clade emerged early in the eudicot lineage—most likely tens of millions of years after the emergence of ZAR1—and has diversified into at least 129 genes in 55 species (Figure 10). ZAR1-SUB proteins are distinctly more diverse in sequence than ZAR1 orthologs and generally lack key sequence features of ZAR1, like the MADA motif and the NBD-NBD oligomerisation interface (Figure 9) (Adachi et al., 2019b; Wang et al. 2019b; Hu et al. 2020). This pattern is consistent with ‘use-it-or-lose-it’ evolutionary model, in which NLRs that specialize for pathogen detection lose some of the molecular features of their multifunctional ancestors (Adachi et al., 2019b). Therefore, we predict that many ZAR1-SUB proteins evolved into specialized sensor NLRs that require NLR helper mates for executing the hypersensitive response. It is possible that ZAR1-SUB helper mate is ZAR1 itself, and that these NLRs evolved into a phylogenetically linked network of sensors and helpers similar to the NRC network of asterid plants (Wu et al., 2017). However, 11 species have a ZAR1-SUB gene but lack a canonical ZAR1 (Supplemental Data Set 2), indicating that these ZAR1-SUB NLRs may have evolved to depend on other classes of NLR helpers.

How would ZAR1-SUB sense pathogens? Given that the LRR domains of most ZAR1-SUB proteins markedly diverged from the RLCK binding interfaces of ZAR1, it is unlikely that ZAR1-SUB proteins bind RLCKs in a ZAR1-type manner (Supplemental Figure 13). This leads us to draw the hypothesis that ZAR1-SUB proteins have diversified to recognize other ligands than RLCKs. In the future, functional investigations of ZAR1-SUB proteins could provide insights into how multifunctional NLRs, such as ZAR1, evolve into functionally specialized NLRs.

The ZAR1-CIN clade consists of 11 clustered paralogs that are unique to the magnoliid species stout camphor as revealed from the genome sequence of the Taiwanese small-flowered camphor tree (also known as *Cinnamomum kanehirae*, Chinese name niu zhang 牛樟) (Chaw et al., 2019). This cluster probably expanded from ZAR1, which is ∼48 Mbp on the same genome sequence scaffold (Supplemental Figure 14). The relatively rapid expansion pattern of ZAR1-CIN into a tandemly duplicated gene cluster is more in line with the classical model of NLR evolution compared to ZAR1 maintenance as a genetic singleton over tens of millions of years (Michelmore and Meyers, 1998). ZAR1-CIN proteins may have neofunctionalized after duplication, acquiring new recognition specificities as consequence of coevolution with host partner proteins and/or pathogen effectors. Consistent with this view, ZAR1-CIN exhibit different patterns of gene expression across tissues (Supplemental Figure 14). Moreover, ZAR1-CIN proteins display distinct surfaces at the ZAR1-RLCK binding interfaces and may bind to other ligands than RLCKs as we hypothesized above for ZAR1-SUB (Supplemental Figure 13). ZAR1-CIN could be viewed as intraspecific highly variable NLRs (hvNLR) per the nomenclature of Prigozhin and Krasileva (2020).

Unlike ZAR1-SUB, ZAR1-CIN have retained the N-terminal MADA sequence (Figure 9, Supplemental Figure 13). We propose that ZAR1-CIN are able to execute the hypersensitive cell death on their own similar to ZAR1. However, ZAR1-CIN display divergent sequence patterns at NBD-NBD oligomerisation interfaces compared to ZAR1 (Figure 9C, Supplemental Figure 15). Therefore, ZAR1-CIN may form resistosome-type complexes that are independent of ZAR1. One intriguing hypothesis is that ZAR1-CIN may associate with each other to form heterocomplexes of varying complexity and functionality operating as an NLR receptor network. In any case, the clear-cut evolutionary trajectory from ZAR1 to the ZAR1-CIN paralog cluster provides a robust evolutionary framework to study functional transitions and diversifications in this CC-NLR lineage.

In summary, our phylogenomics analyses raise several intriguing questions about ZAR1 evolution. The primary conclusion we draw is that ZAR1 is an ancient CC-NLR that has been a partner with RLCKs ever since the Jurassic Period. We propose that throughout at least 150 million years, ZAR1 has maintained its molecular features for sensing pathogens via RLCKs and activating hypersensitive cell death. Further comparative analyses, combining molecular evolution and structural biology, of plant resistosomes and between resistosomes and the apoptosomes and inflammasome of animal NLR systems (Wang and Chai, 2020) will yield novel experimentally testable hypotheses for NLR research.

## Materials and Methods

### ZAR1 and ZRK sequence retrieval

We performed BLAST (Altschul et al., 1990) using previously identified ZAR1 and ZRK sequences as queries (Lewis et al., 2013; Baudin et al., 2017; Schultink et al., 2019; Harant et al., 2021) to search ZAR1 and ZRK like sequences in NCBI nr or nr/nt database (https://blast.ncbi.nlm.nih.gov/Blast.cgi) and Phytozome12.1 (https://phytozome.jgi.doe.gov/pz/portal.html#!search?show=BLAST). In the BLAST search, we used cut-offs, percent identity ≥ 30% and query coverage ≥ 80%. The BLAST pipeline was circulated by using the obtained sequences as new queries to search ZAR1 and ZRK like genes over the angiosperm species. We also performed the BLAST pipeline against a plant NLR dataset annotated by NLR-parser (Steuernagel et al., 2015) from 38 plant reference genome databases (Supplemental Data Set 11).

### Phylogenetic analyses

For the phylogenetic analysis, we aligned NLR and ZRK amino acid sequences (Supplemental Data Sets 5, 12 to 16) using MAFFT v.7 (Katoh and Standley, 2013) and manually deleted the gaps in the alignments in MEGA7 (Kumar et al., 2016). Full-length or NB-ARC domain sequences of the aligned NLR datasets were used for generating phylogenetic trees. To generate ZRK phylogenetic trees, we used full-length or kinase domain sequences of the aligned ZRK datasets. The neighbour-joining tree was made using MEGA7 with JTT model and bootstrap values based on 100 iterations. All phylogenetic tree files are in Supplemental Data Sets 17 to 21.

### Patristic distance analyses

To calculate the phylogenetic (patristic) distance, we used Python script based on DendroPy (Sukumaran and Mark, 2010). We calculated patristic distances from each CC-NLR to the other CC-NLRs on the phylogenetic tree and extracted the distance between CC-NLRs of Arabidopsis or *N. benthamiana* to the closest NLR from the other plant species. The script used for the patristic distance calculation is available from GitHub (https://github.com/slt666666/Phylogenetic_distance_plot2).

### Gene co-linearity analyses

To investigate genetic co-linearity at ZAR1 loci, we extracted the 3 genes upstream and downstream of ZAR1 using GFF files derived from reference genome databases (Supplemental Data Set 11). To identify conserved gene blocks, we used gene annotation from NCBI Protein database and confirmed protein domain information based on InterProScan (Jones et al, 2014).

### Sequence conservation analyses

Full-length NLR sequences of each subfamily ZAR1, ZAR1-SUB or ZAR1-CIN were subjected to motif searches using the MEME (Multiple EM for Motif Elicitation) (Bailey and Elkan, 1994) with parameters ‘zero or one occurrence per sequence, top twenty motifs’, to detect consensus motifs conserved in ≥90% of the input sequences. The output data are summarized in Supplemental Tables 1, 2 and 4.

To predict the MADA motif from ZAR1, ZAR1-SUB and ZAR1-CIN datasets, we used the MADA-HMM previously developed (Adachi et al., 2019b), with the hmmsearch program (hmmsearch –max -o <outputfile> <hmmfile> <seqdb>) implemented in HMMER v2.3.2 (Eddy, 1998). We termed sequences over the HMMER cut-off score of 10.0 as the MADA motif and sequences having the score 0-to-10.0 as the MADA-like motif.

To analyze sequence conservation and variation in ZAR1, ZAR1-SUB and ZAR1-CIN proteins, aligned full-length NLR sequences (Supplemental Data Sets 10, 22, 23) were used for ConSurf (Ashkenazy et al., 2016). Arabidopsis ZAR1 (NP_190664.1), a tomato ZAR1-SUB (XP_004243429.1) or a Stout camphor ZAR1-CIN (RWR85656.1) was used as a query for each analysis of ZAR1, ZAR1-SUB or ZAR1-CIN, respectively. The output datasets are in Supplemental Data Sets 24 to 26.

### Protein structure analyses

The atomic coordinate of ZAR1 (protein data bank accession codes; 6J5T) was downloaded from protein data bank for illustration in ccp4mg. We used the cryo-EM structures of ZAR1 as templates to generate homology models of ZAR1-SUB and ZAR1-CIN. Amino acid sequences of a tomato ZAR1-SUB (XP_004243429.1) and a stout camphor ZAR1-CIN (RWR85656.1) were submitted to Protein Homology Recognition Engine V2.0 (Phyre2) for modelling (Kelley et a., 2015). The coordinates of ZAR1 structure (6J5T) were retrieved from the Protein Data Bank and assigned as modelling template by using Phyre2 Expert Mode. The resulting models of ZAR1-SUB and ZAR1-CIN, and the ZAR1 structures (6J5T) were illustrated with the ConSurf conservation scores in PyMol.

### Plant growth condition

Wild-type *N. benthamiana* plants were grown in a controlled growth chamber with temperature 22-25 °C, humidity 45-65% and 16/8 hr light/dark cycle.

### Plasmid constructions

The Golden Gate Modular Cloning (MoClo) kit (Weber et al., 2011) and the MoClo plant parts kit (Engler et al., 2014) were used for cloning, and all vectors are from this kit unless specified otherwise. ZAR1 and RLCK homologs identified in the taro (*Colocasia esculenta*; Assembly: ASM944546v1), columbine (*Aquilegia coerulea*; Assembly: Aquilegia_coerulea_v1; Filiault et al., 2018), and stout camphor tree (*Cinnamomum kanehirae*; Assembly: ASBRC_Ckan_1.0; Chaw et al., 2019) genomes were codon-optimized for *N. benthamiana* using the ThermoFisher GeneOptimizer tool and synthesized by GENEWIZ as Golden Gate Level 0 modules into pICH41155. Genes were subcloned into the binary vector pICH86988 (Weber et al., 2011) and transformed into *A. tumefaciens* strain GV3101 pMP90. Cloning design and sequence analysis were done using Geneious Prime (v2022.0.1; https://www.geneious.com). Plasmid construction is described in Supplemental Data Set 27.

### Transient gene-expression and cell death assay

Transient expression of NbZAR1 mutants in *N. benthamiana* were performed by agroinfiltration according to methods described by Bos et al. (2006). Briefly, four-weeks old *N. benthamiana* plants were infiltrated with *Agrobacterium tumefaciens* strains carrying the binary expression plasmids. *A. tumefaciens* suspensions were prepared in infiltration buffer (10 mM MES, 10 mM MgCl_2_, and 150 μM acetosyringone, pH5.6) and were adjusted to appropriate OD_600_ (Supplemental Table 13). Macroscopic cell death phenotypes were scored according to the scale of Segretin et al. (2014) modified to range from 0 (no visible necrosis) to 7 (fully confluent necrosis).

### RNA-seq data analyses

Public RNA-seq reads, which were previously obtained with Illumina HiSeq 2000 (Chaw et al., 2019), were used to analyze expression profiles of CmZAR1 and ZAR1-CIN genes in the stout camphor tree (Accession Numbers: SRR7416905, SRR7416906, SRR7416908, SRR7416909, SRR7416910, SRR7416911, and SRR7416918). Reads were mapped to the stout camphor genome assembly (GenBank assembly accession GCA_003546025.1) using the splice-aware RNAseq tool in CLC Genomics Workbench vs 20.0.4 (https://digitalinsights.qiagen.com) and transformed into a Transcripts Per Million (TPM) value according to Li et al. (2010). TPM values were visualized by the heatmap. The heatmap was colored by eight ranges (0, 0∼5, 5∼20, 20∼40, 40∼60, 60∼80, 80∼100, 100<) of TPM values.

## Accession numbers

DNA sequence data used in this study can be found from reference genome or GenBank/EMBL databases with accession numbers listed in Supplemental Data Sets 1, 4, 8 and 9.

## Supplemental Data

**Supplemental Figure 1.**
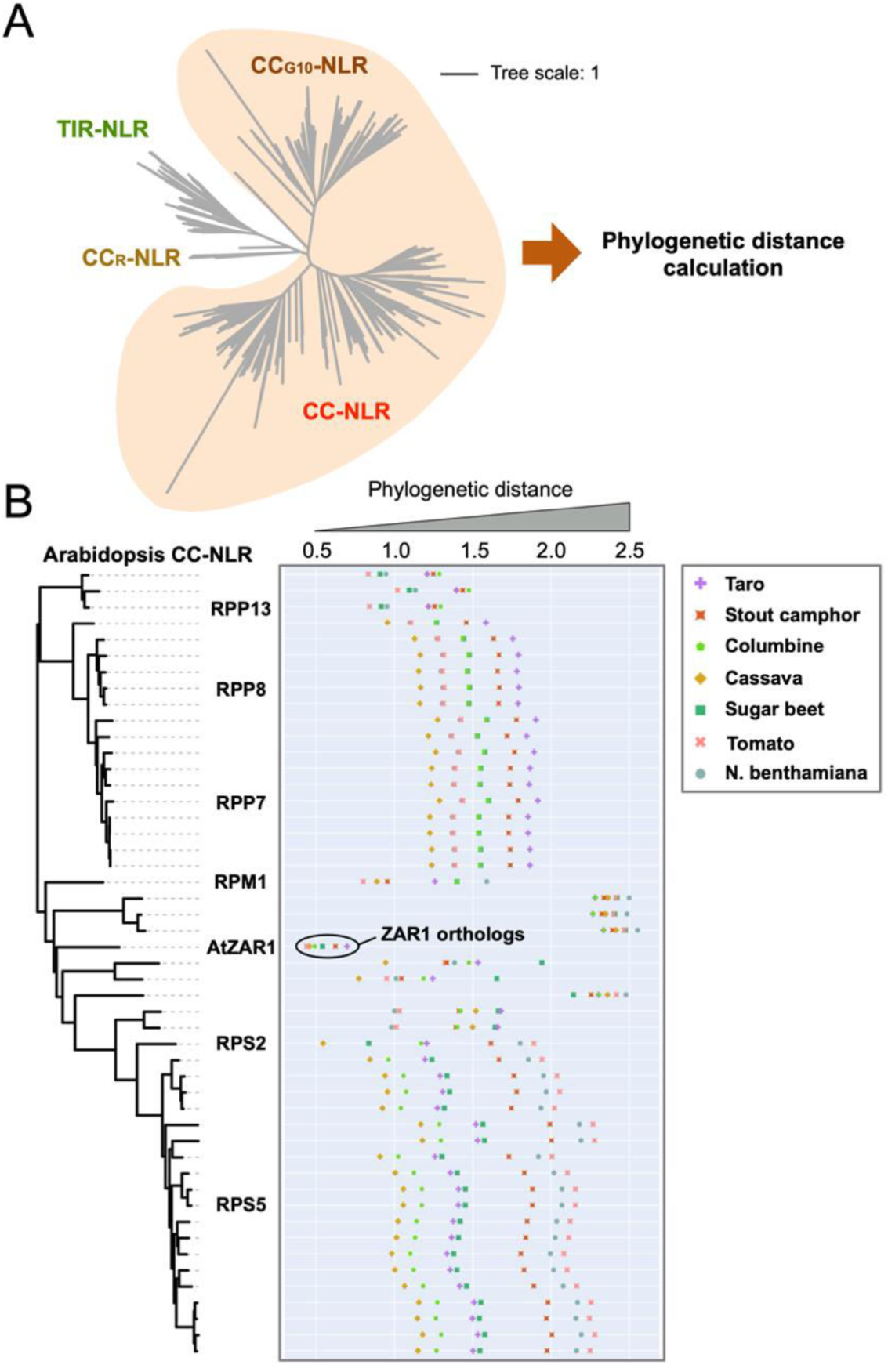
Arabidopsis ZAR1 is the most conserved CC-NLR across angiosperms. (**A**) Phylogenetic tree of NLR proteins from 8 plant species. The phylogenetic tree was generated in MEGA7 by the neighbour-joining method using NB-ARC domain sequences of 1475 NLRs identified from taro, stout camphor, columbine, Arabidopsis, cassava, sugar beet, tomato and *N. benthamiana*. The scale bars indicate the evolutionary distance in amino acid substitution per site. We used CC-NLR and CCG10-NLR superclades for calculating phylogenetic distances. (**B**) The phylogenetic (patristic) distance of two CC-NLR nodes between Arabidopsis and other plant species were calculated from the NB-ARC phylogenetic tree in A. The closest patristic distances are plotted with different colours based on plant species. Representative Arabidopsis NLRs are highlighted. The closest patristic distances of two CC-NLR nodes between *N. benthamiana* and other plant species can be found in Supplemental Figure 2.

**Supplemental Figure 2.**
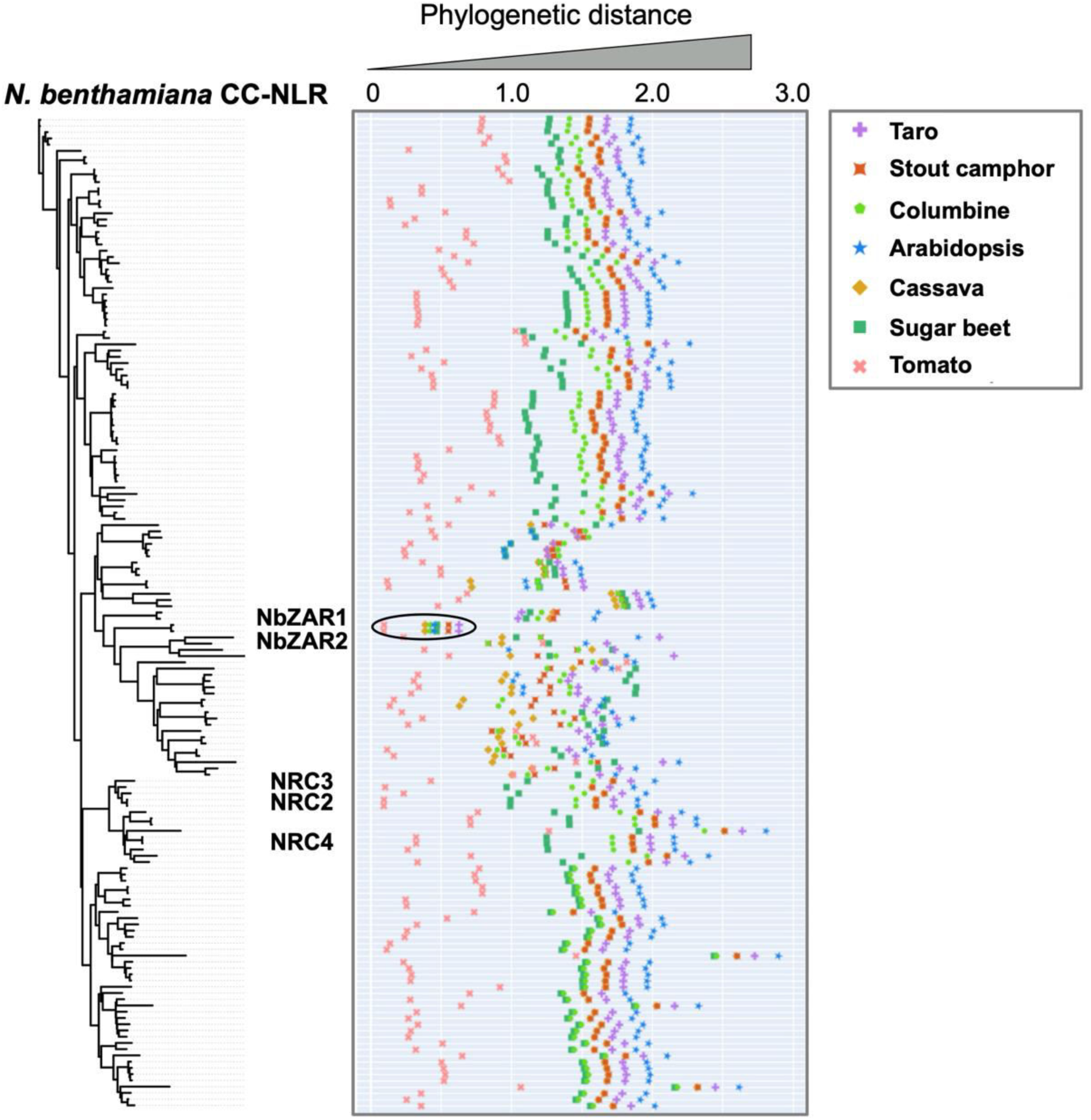
NbZAR1 is highly conserved across angiosperms. The phylogenetic (patristic) distance of two CC-NLR nodes between *N. benthamiana* and the closest NLR from the other plant species were calculated from the NB-ARC phylogenetic tree in Supplemental Figure 1. The closest patristic distances are plotted with different colours based on plant species.

**Supplemental Figure 3.**
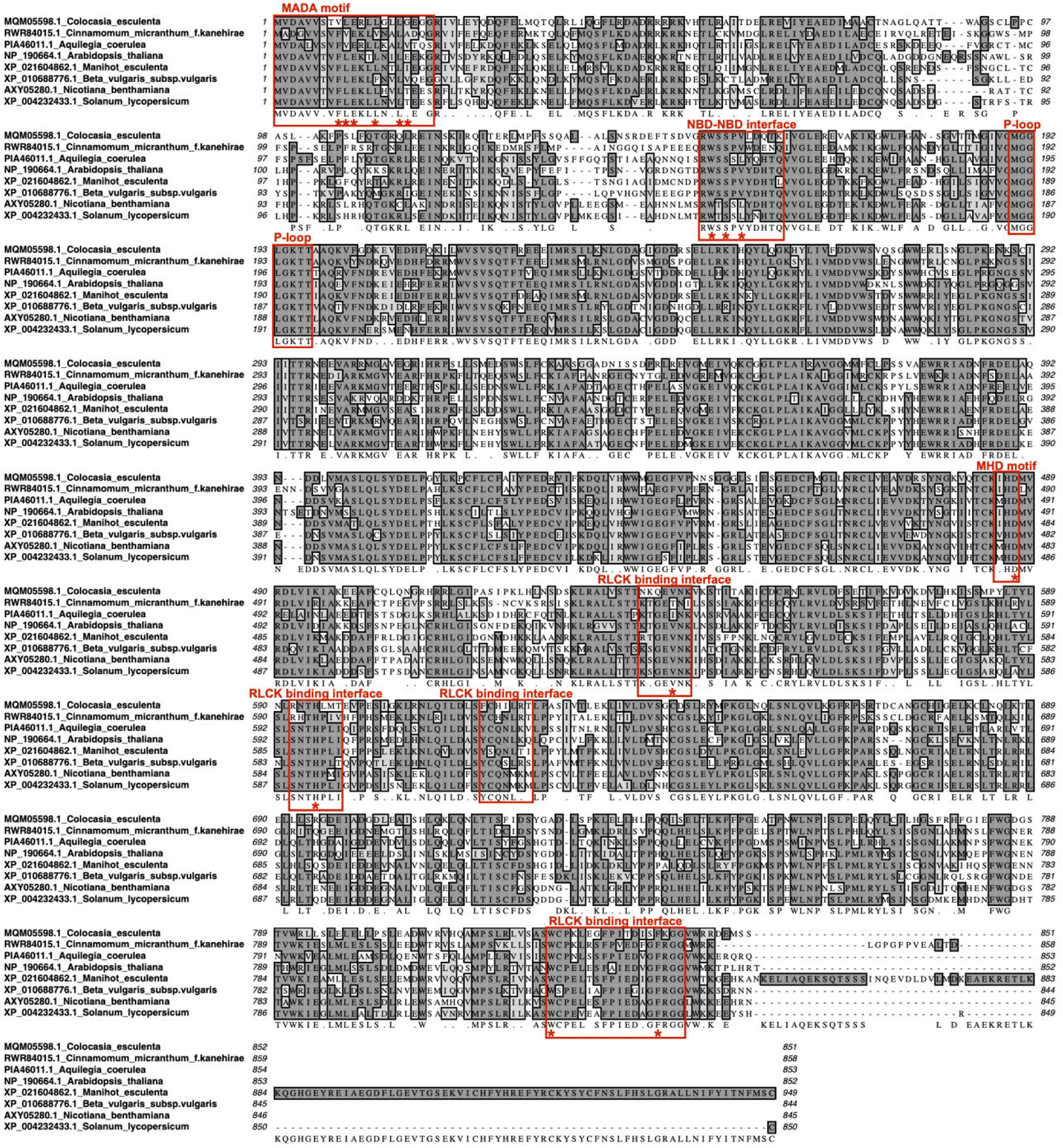
Sequence alignment of full-length ZAR1 ortholog proteins across angiosperms. Amino acid sequences of ZAR1 orthologs were aligned by MAFFT version 7 program. Conserved motif sequences highlighted in this study are marked with red boxes. Red asterisks indicate substitution sites for introducing gain or loss of ZAR1 protein function.

**Supplemental Figure 4.**
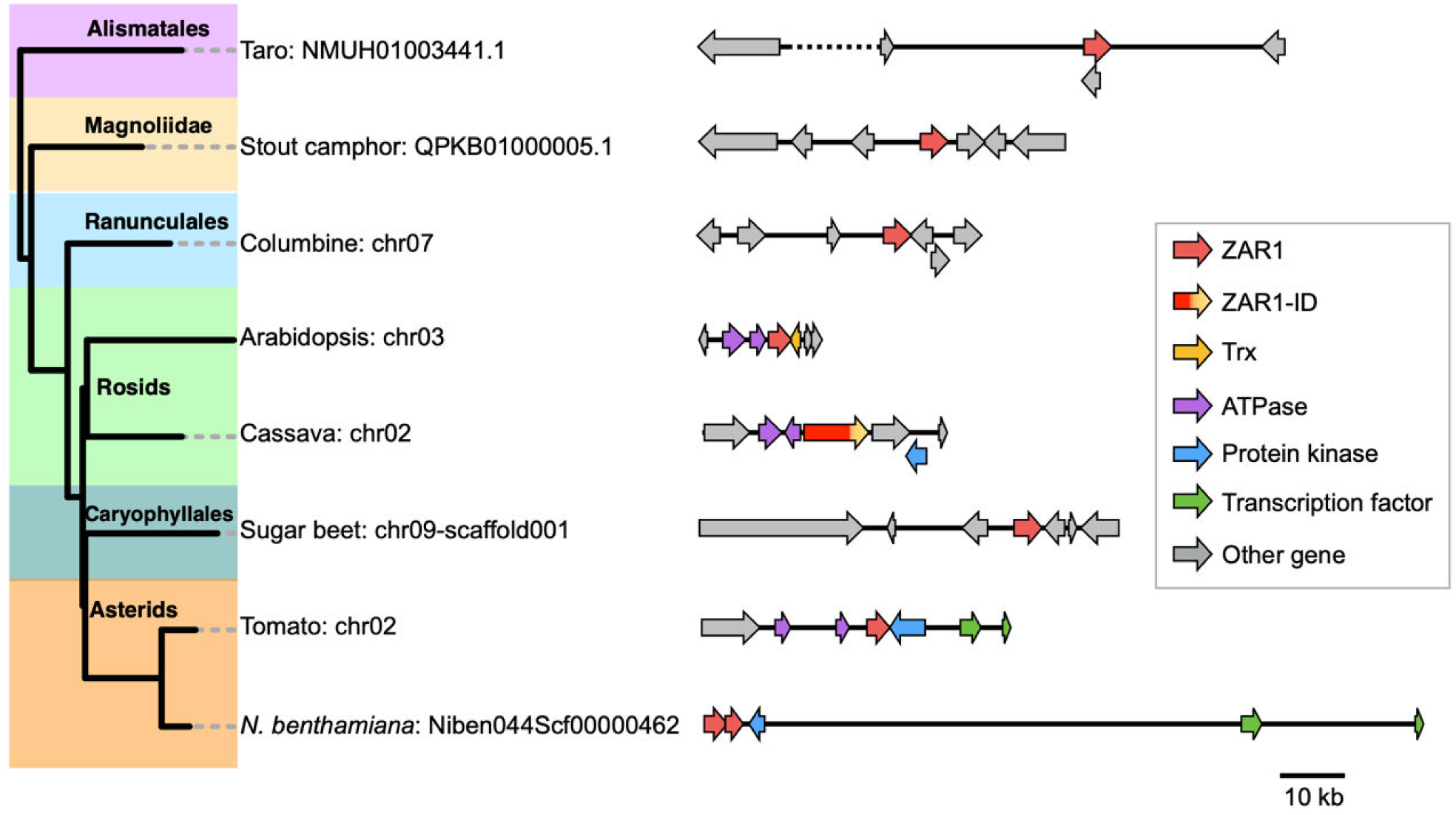
Schematic representation of the intragenomic relationship at ZAR1 loci across angiosperm genomes. We selected representative 8 plant species genome assemblies based on the phylogenetic tree in Figure 2 and used them for the synteny-based analysis of the ZAR1 loci. We highlight genes showing intragenomic linkages with different colours based on the gene annotations. Genes genetically linked to ZAR1 in eudicots are listed in Supplemental Data Set 7.

**Supplemental Figure 5.**
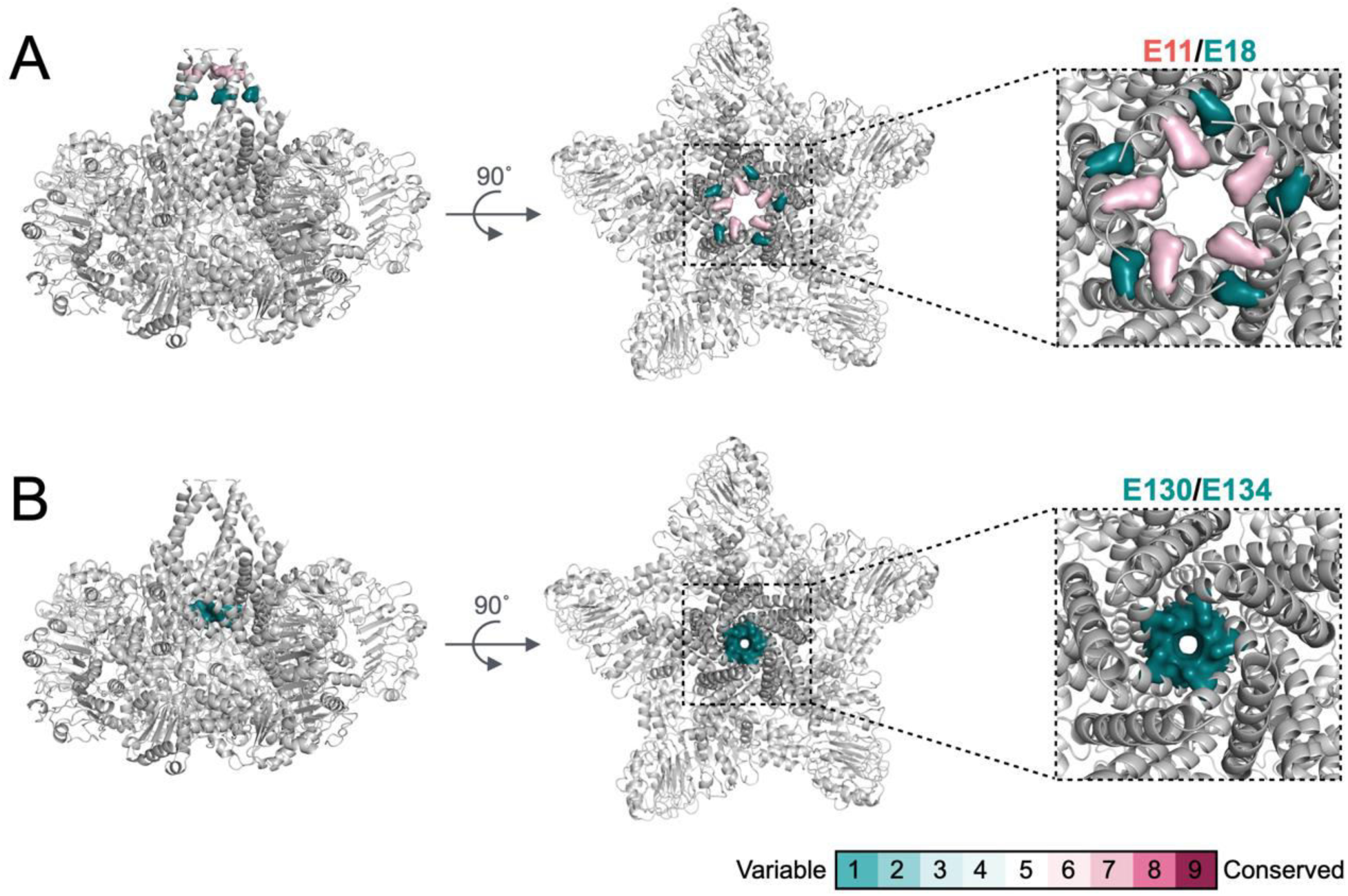
E11 on glutamate ring inside of the Arabidopsis ZAR1 resistosome is conserved across the orthologs. The ConSurf conservation scores at E11 and E18 (**A**) or at E130 and E134 (**B**) are illustrated in cartoon representation of the Arabidopsis ZAR1 resistosome structure.

**Supplemental Figure 6.**
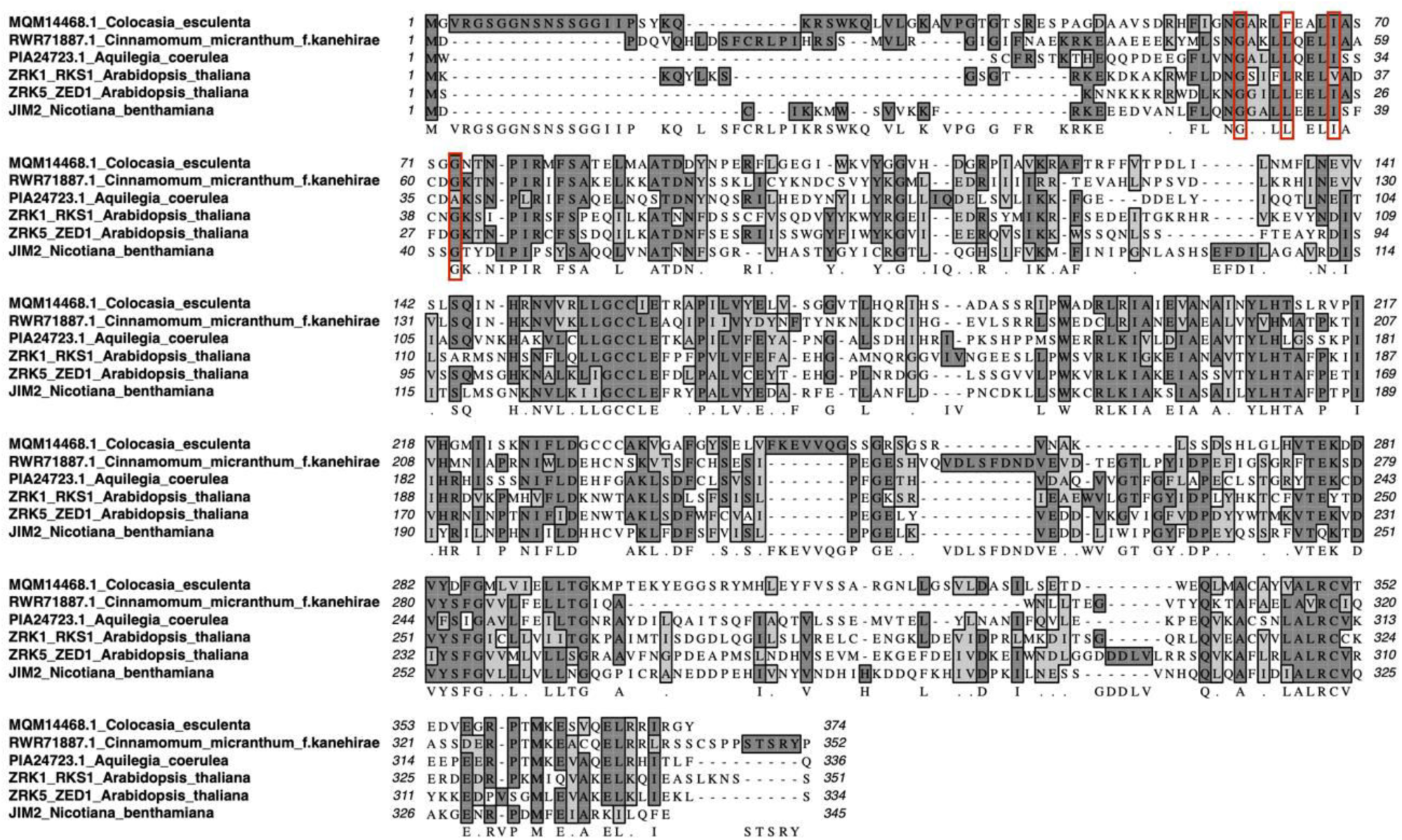
Sequence alignment of full-length ZRK proteins across angiosperms. Amino acid sequences of ZRK proteins were aligned by MAFFT version 7 program. Functionally validated residues for ZRK-ZAR1 interactions are marked with red boxes.

**Supplemental Figure 7.**
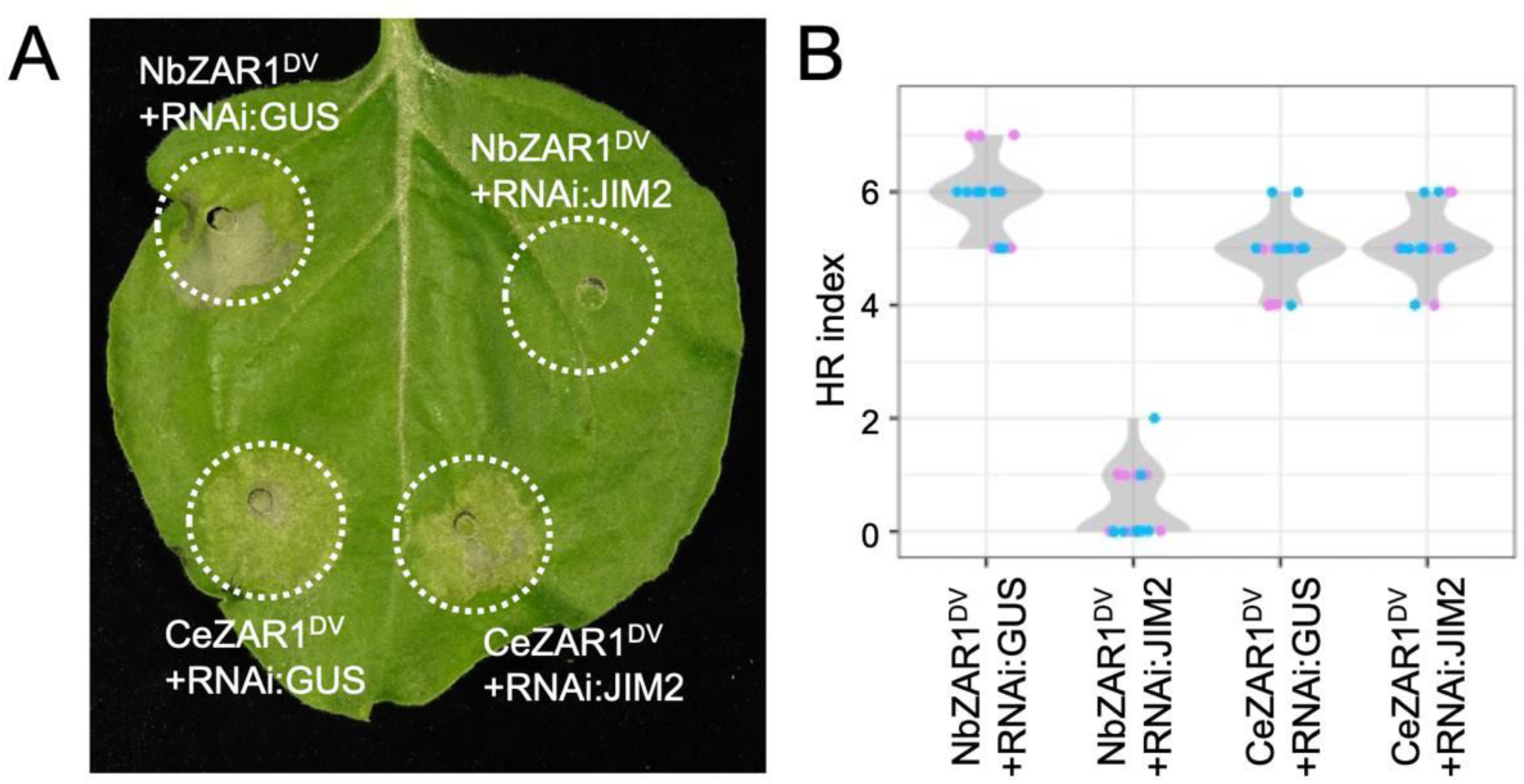
Silencing of JIM2 does not affect CeZAR1 autoactive cell death in *N. benthamiana*. (**A**) Cell death observed in *N. benthamiana* leaves after expression of autoactive NbZAR1 (NbZAR1^DV^) and CeZAR1 (CeZAR1^DV^) with RNAi constructs. *N. benthamiana* leaf panels were photographed at 4 to 6 days after agroinfiltration. (**B**) Violin plots showing cell death intensity scored as an HR index based on 18 replicates from two independent experiments.

**Supplemental Figure 8.**
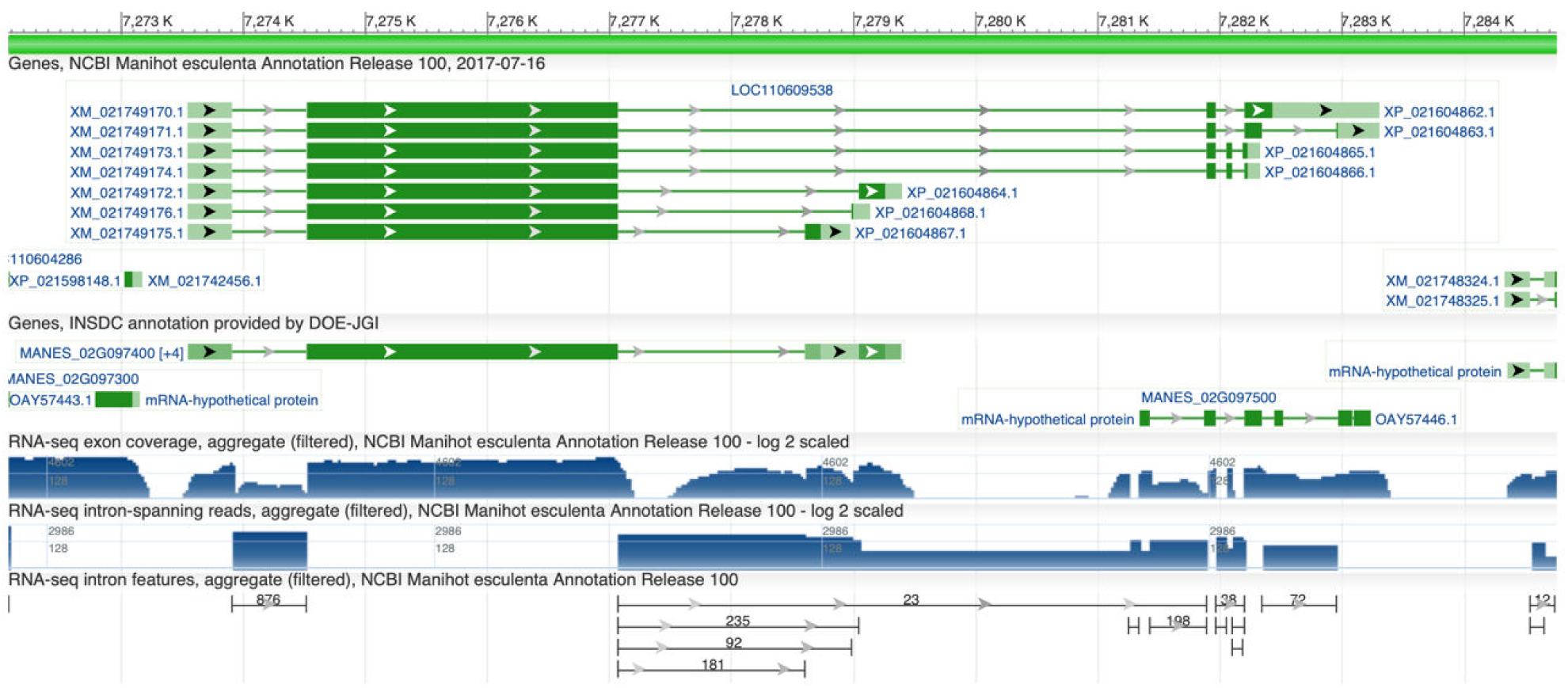
Cassava ZAR1 and ZAR1-ID are transcribed from a single locus on the genome. The gene locus of cassava ZAR1 (XP_021604863.1, XP_021604865.1, XP_021604866.1, XP_021604867.1 and XP_021604868.1) and ZAR1-ID (XP_021604862.1 and XP_021604864.1) is shown with RNA-seq exon coverage and is extracted from NCBI database (database ID: LOC110609538).

**Supplemental Figure 9.**
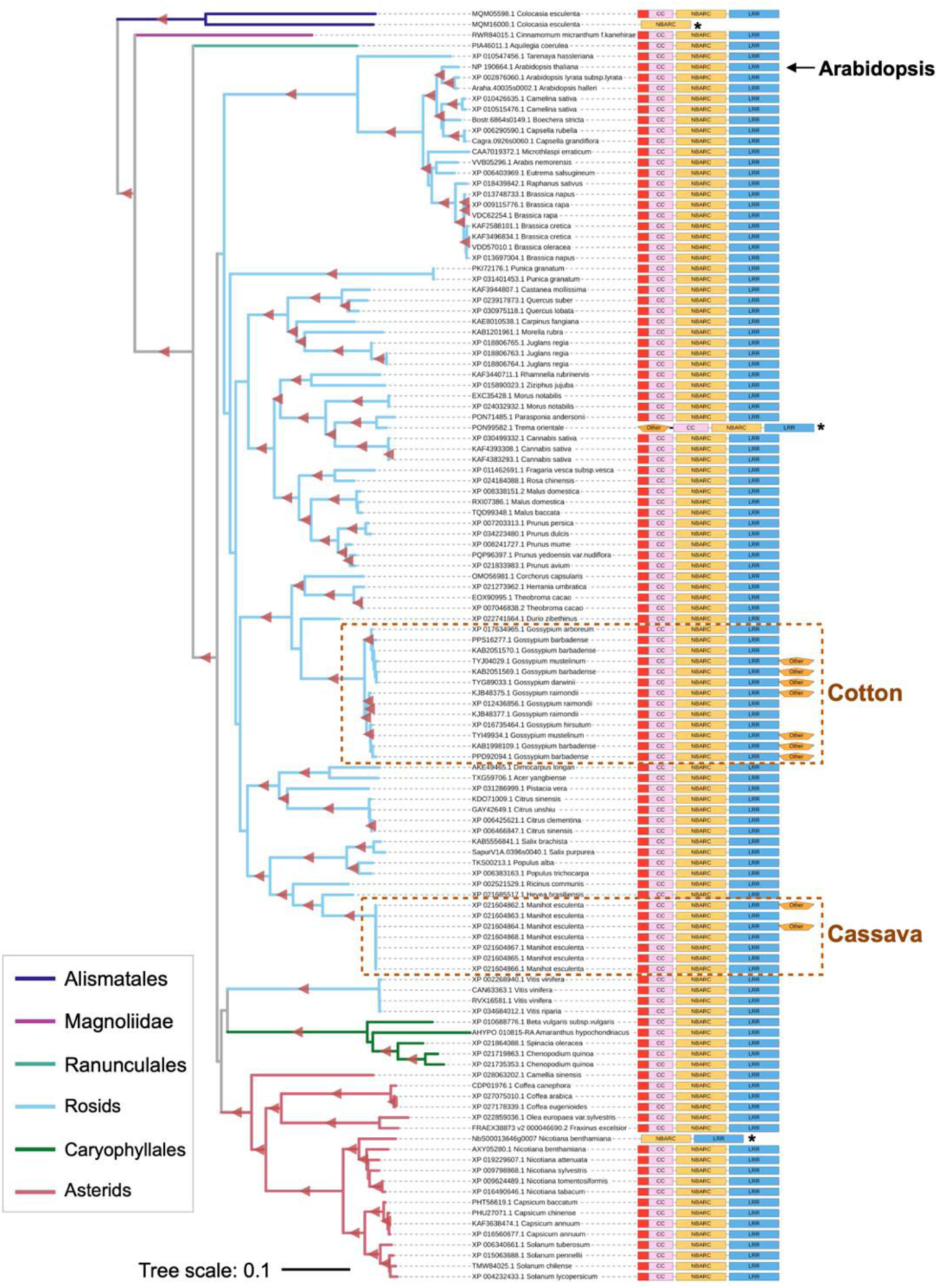
Trx domain integration occurred in two independent rosid ZAR1 subclades. The phylogenetic tree shown in Figure 2 was used to describe NLR domain architectures. Domain schemes are aligned to right side of the leaf labels: MADA is red, CC is pink, NB-ARC is yellow, LRR is blue and other domain is orange. Black asterisks on domain schemes describe truncated NLRs or potentially mis-annotated NLR. Each branch is marked with different colours based on the plant taxonomy. Red triangles indicate bootstrap support > 0.7. The scale bar indicates the evolutionary distance in amino acid substitution per site.

**Supplemental Figure 10.**
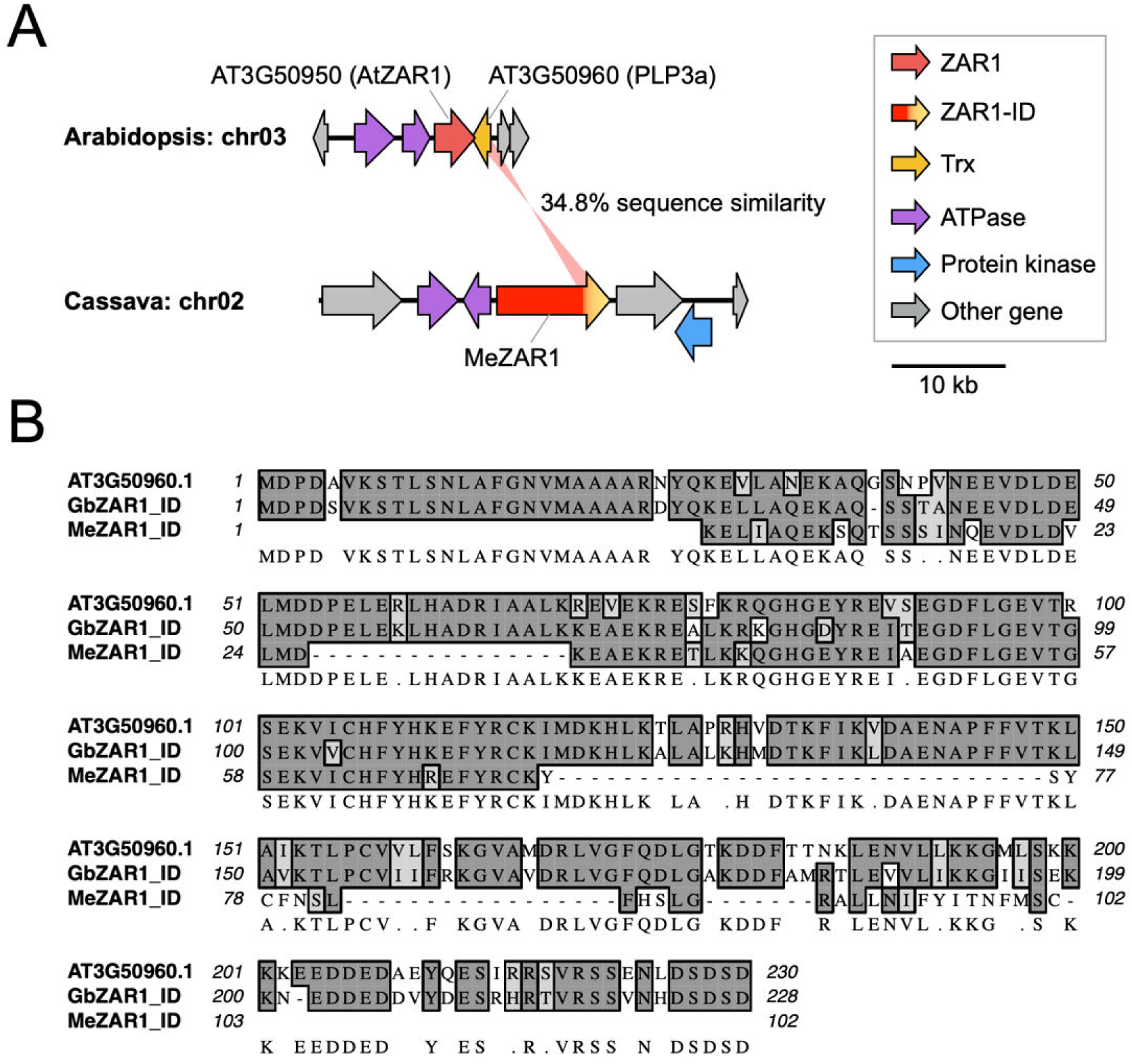
Integrated Trx domains show high sequence similarity to ZAR1-linked PLP3a gene in Arabidopsis. (**A**) Schematic representation of the intragenomic relationship at ZAR1 loci between Arabidopsis and cassava. We highlight sequence similarity of integrated Trx domain in Cassava ZAR1 (MeZAR1) to PLP3a gene genetically linked to Arabidopsis ZAR1 (AtZAR1). Details are explained in Supplemental Figure 4. (**B**) Amino acid sequences of Arabidopsis PLP3a gene (AT3G50960) and integrated domains of an MeZAR1 (XP_021604862.1) and a cotton ZAR1 (GbZAR1; KAB1998109.1).

**Supplemental Figure 11.**
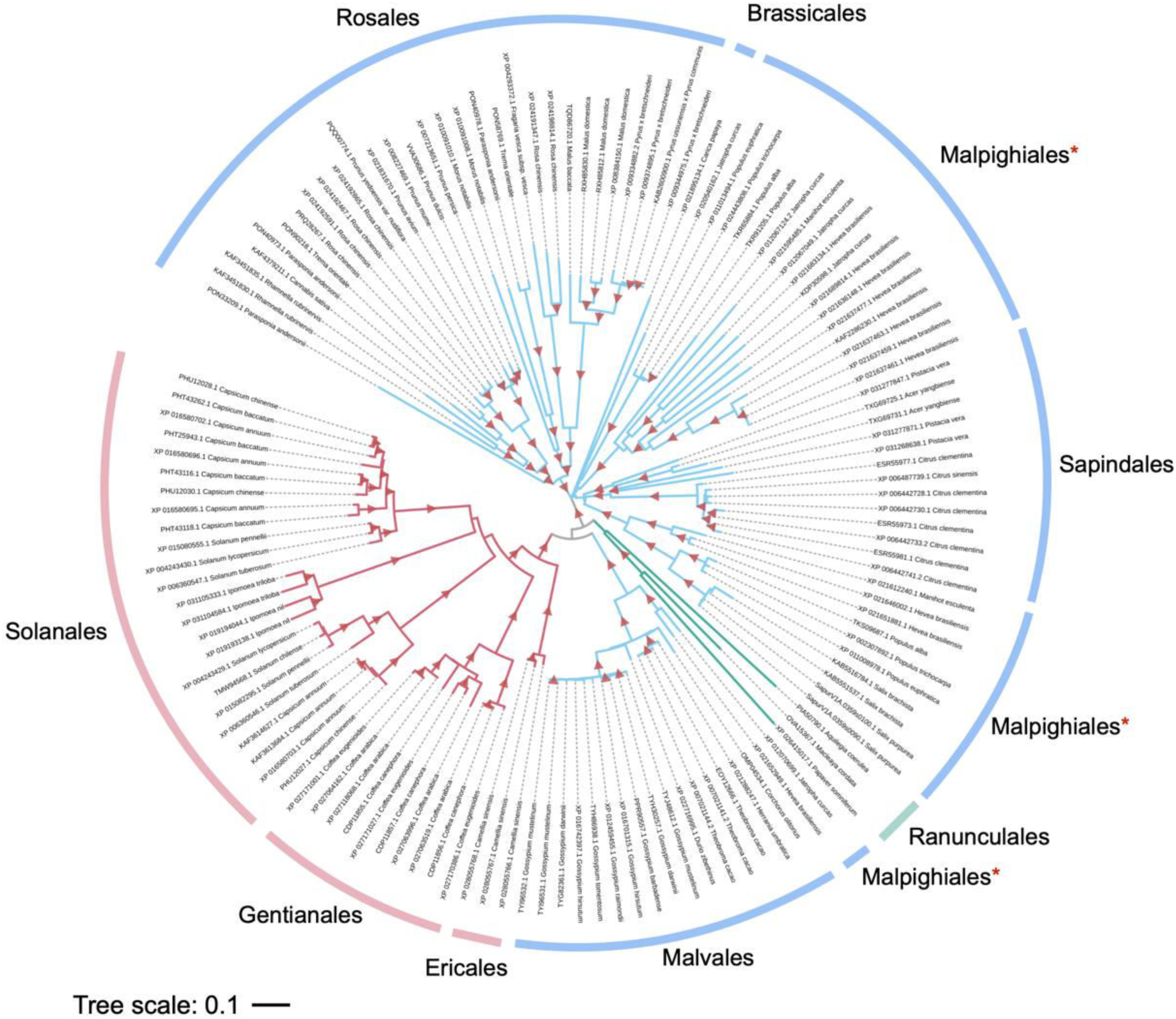
ZAR1-SUB gene is distributed across eudicots. The phylogenetic tree was generated in MEGA7 by the neighbour-joining method using full length amino acid sequences of 129 ZAR1-SUB orthologs identified in Figure 1. Each branch is marked with different colours based on the plant taxonomy. Red triangles indicate bootstrap support > 0.7. The scale bar indicates the evolutionary distance in amino acid substitution per site. Red asterisks on plant order term describe that NLRs from Malpighiales are distributed in three independent clades.

**Supplemental Figure 12.**
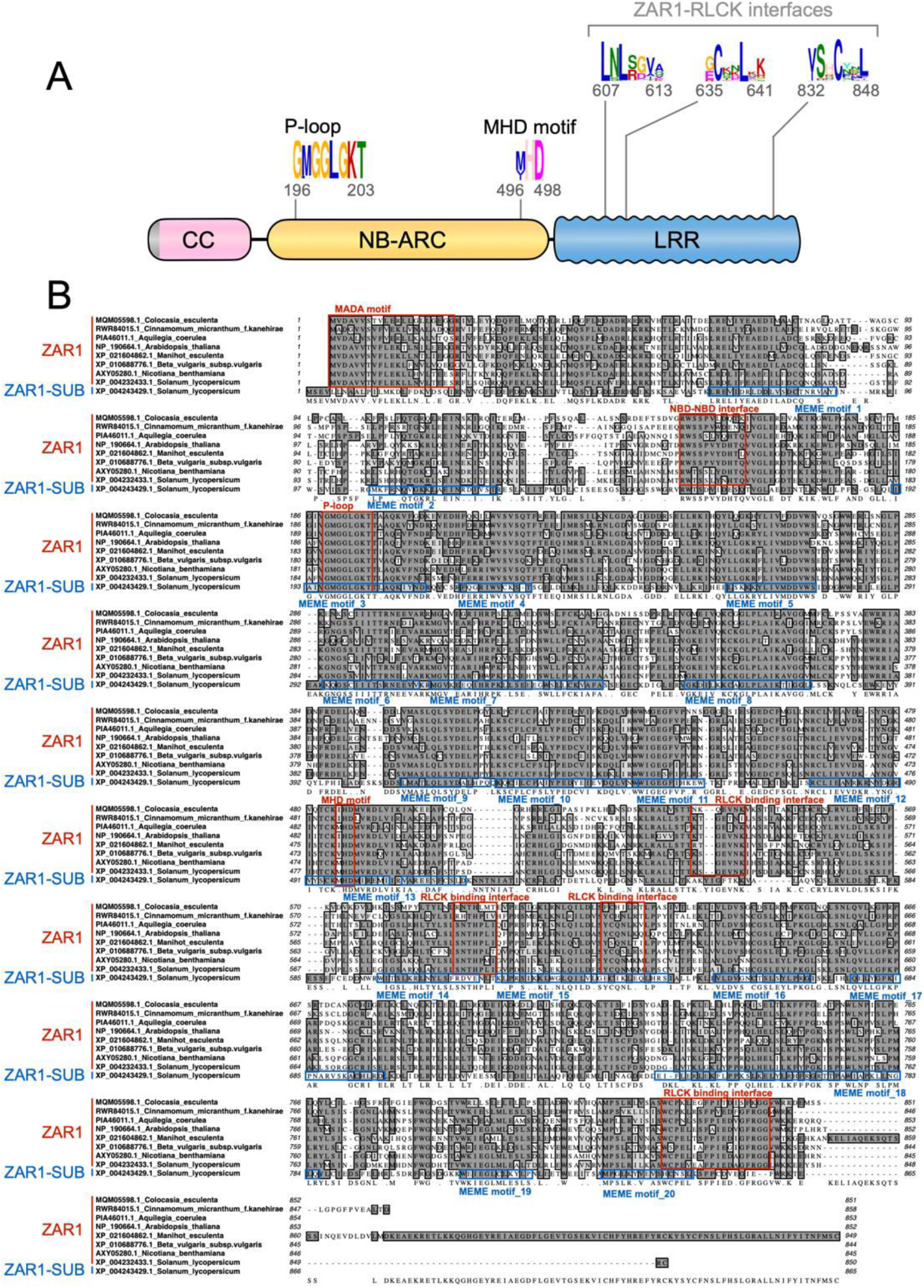
Sequence alignment of full-length ZAR1 and ZAR1-SUB proteins. (**A**) Schematic representation of the ZAR1-SUB protein highlighting the position of the representative conserved sequence patterns across ZAR1-SUB. (**B**) Amino acid sequences of ZAR1 orthologs and a representative ZAR1-SUB (XP_004243429.1 from tomato) were aligned by MAFFT version 7 program. ZAR1 motif sequences highlighted in this study are marked with red boxes. Positions of MEME motifs identified from ZAR1-SUB are marked in blue boxes. Raw MEME motifs are listed in Supplemental Tables 2 and 3.

**Supplemental Figure 13.**
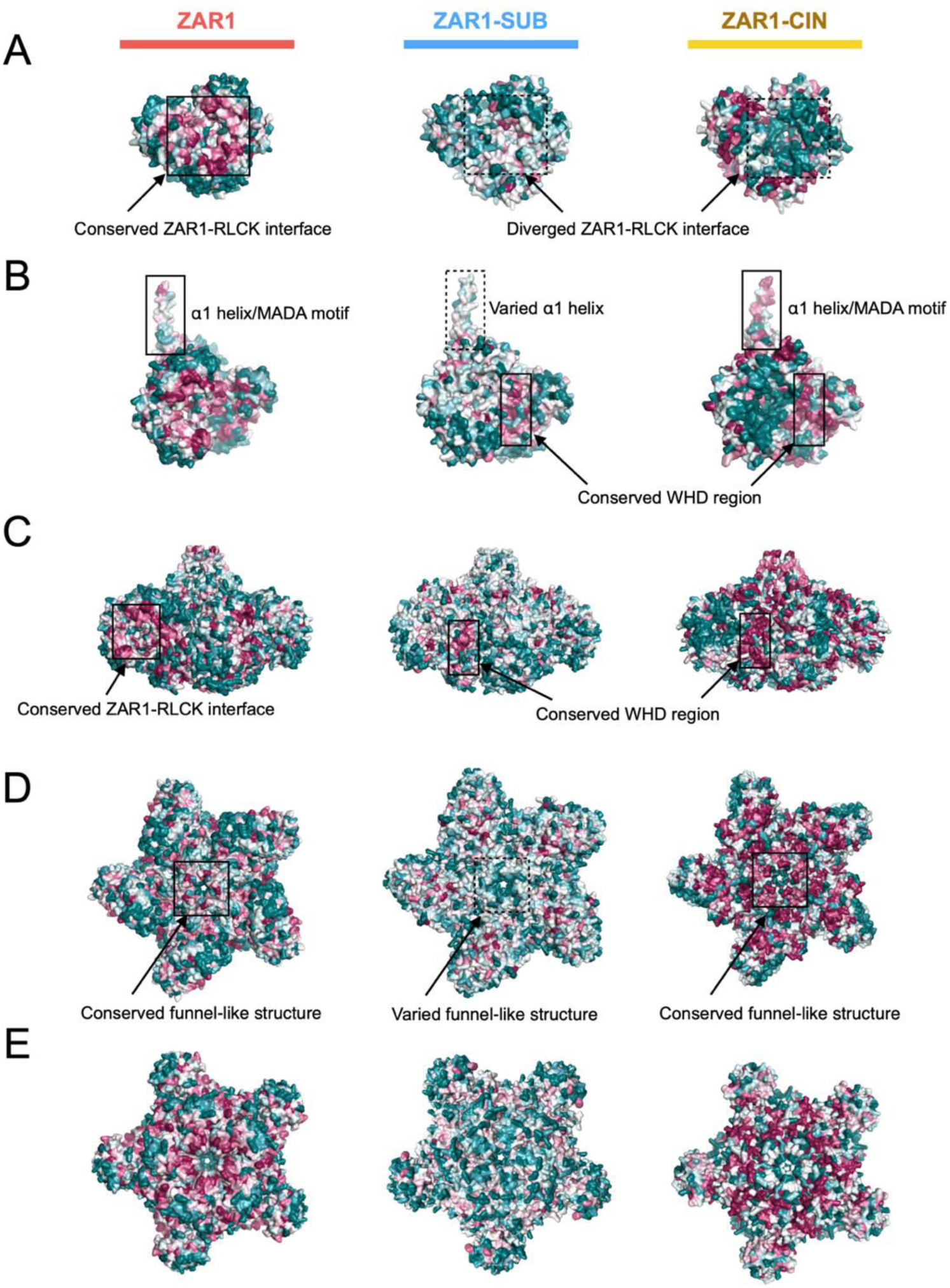
ZAR1 and the sister subclade NLRs display different conserved surfaces on the resistosome structure. Distribution of the ConSurf conservation score was visualized on the inactive monomer (**A**), active monomer (**B**) and resistosome structures (**C-E**) of Arabidopsis ZAR1 or the structure homology models of ZAR1-SUB (XP_004243429.1) and ZAR1-CIN (RWR85656.1). Each structure and cartoon representation are illustrated with different colours based on the conservation score shown in Figures 3 and 9. Resistosome structures are shown from different angles, from side (C), from upper side (**D**) and from underside (E).

**Supplemental Figure 14.**
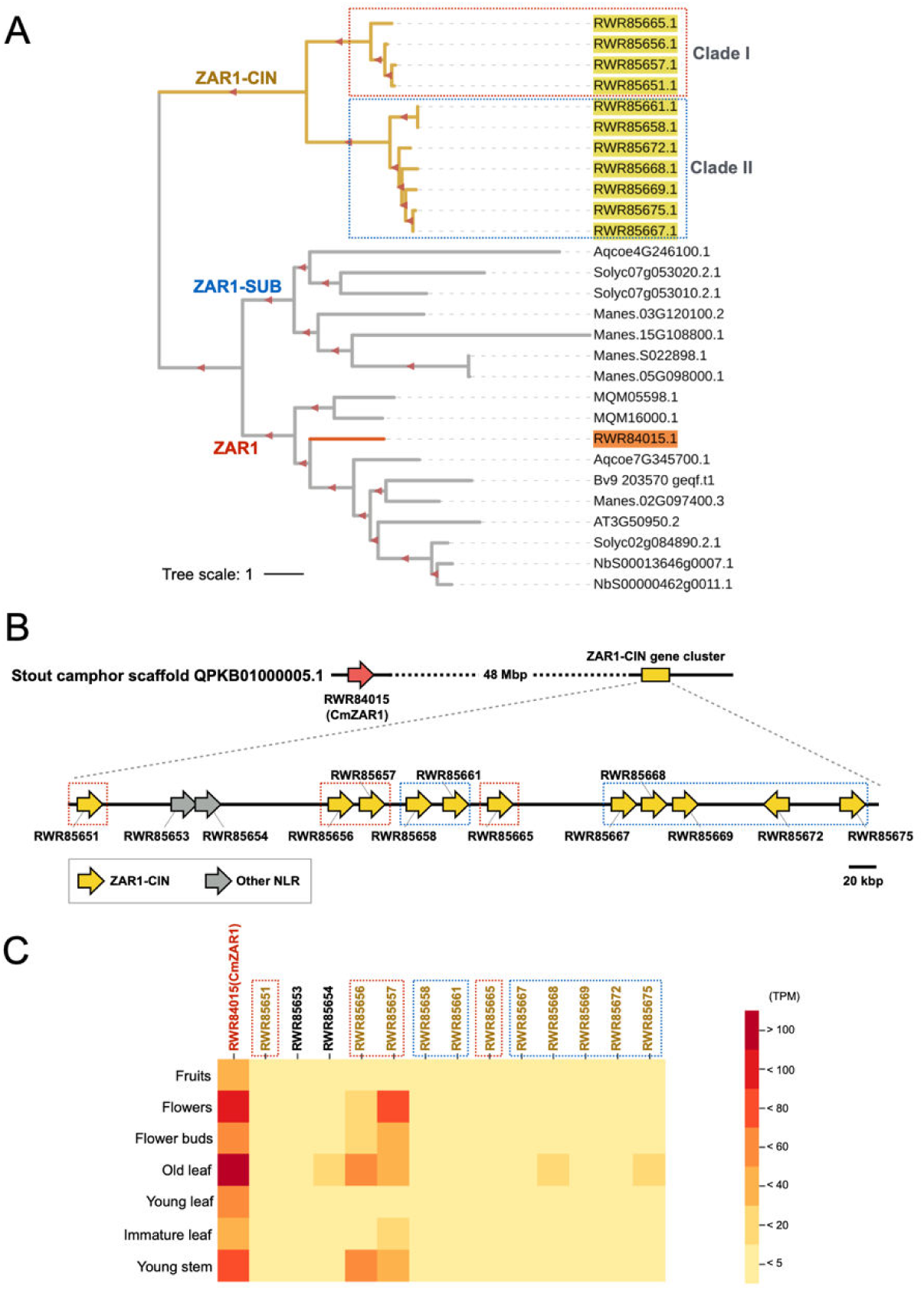
ZAR1-CIN gene cluster occurs in the *Cinnamomum micranthum* genome. (**A**) The subclades including ZAR1, ZAR1-SUB and ZAR1-CIN were zoomed in from the phylogenetic tree constructed in Supplemental Figure 1. Red triangles indicate bootstrap support > 0.7. The scale bar indicates the evolutionary distance in amino acid substitution per site. Well supported subclades (I and II) in ZAR1-CIN are described with red or blue dot box. The gene IDs: taro (MQM-), stout camphor (RWR-), columbine (Aqcoe-), Arabidopsis (AT-), cassava (Manes-), sugar beet (Bv-), tomato (Solyc-) and *N. benthamiana* (NbS-). (**B**) Schematic representation of the ZAR1-CIN gene cluster on a *C. micranthum* (Stout camphor) scaffold. Stout camphor ZAR1 (CmZAR1) and ZAR1-CIN genes are highlighted in orange and yellow, respectively. (**C**) A heatmap showing Transcripts Per Million (TPM) values of the CmZAR1 and ZAR1-CIN genes. Public RNA-seq datasets from seven different tissue samples in *C. micranthum* were used for this heatmap analysis.

**Supplemental Figure 15.**
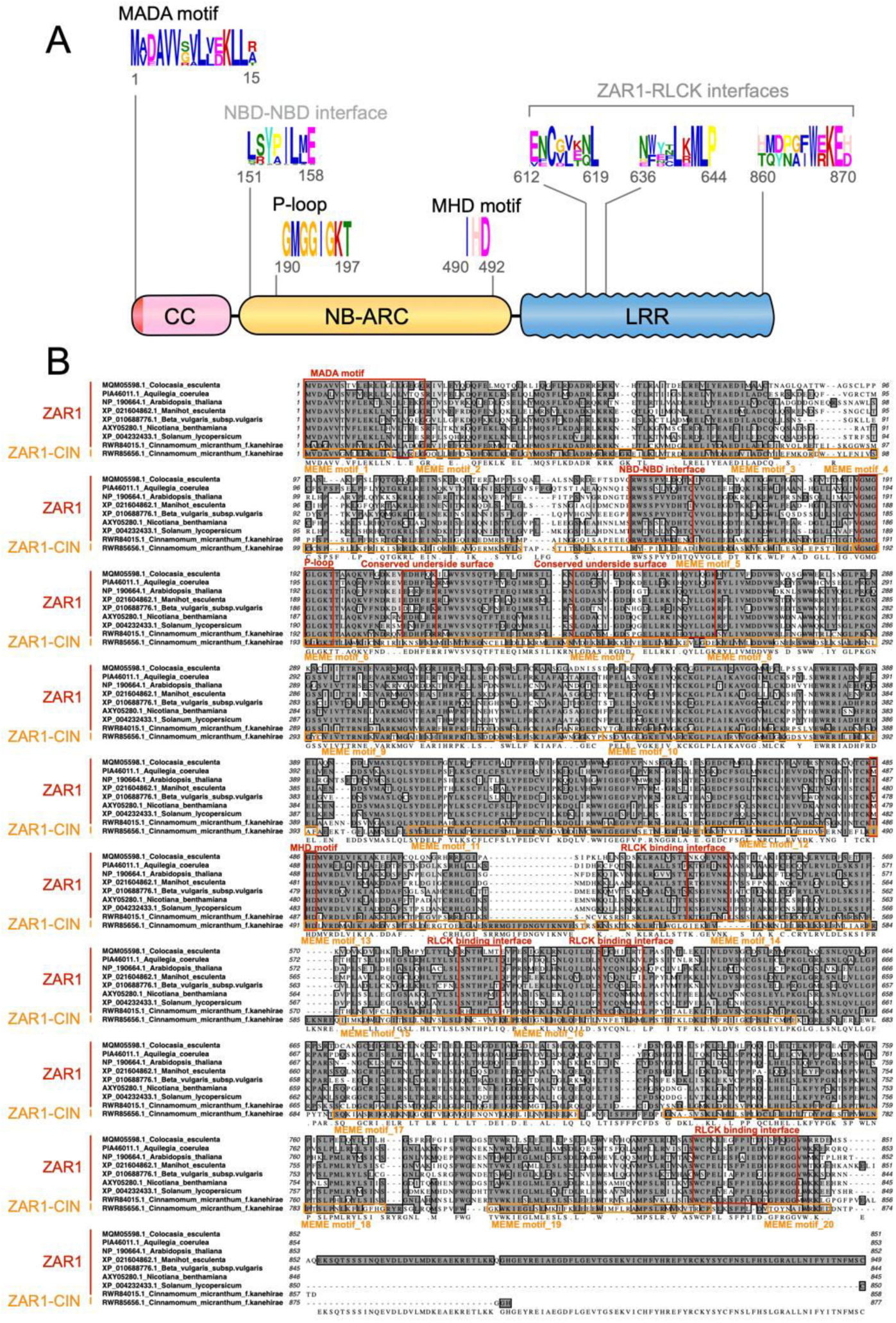
Sequence alignment of full-length ZAR1 and ZAR1-CIN proteins. (**A**) Schematic representation of the ZAR1-CIN protein highlighting the position of the representative conserved sequence patterns across ZAR1-SUB. (**B**) Amino acid sequences of ZAR1 orthologs and a representative ZAR1-CIN (RWR85656.1) were aligned by MAFFT version 7 program. ZAR1 motif sequences highlighted in this study are marked with red boxes. Positions of MEME motifs identified from ZAR1-CIN are marked in orange boxes. Raw MEME motifs are listed in Supplemental Tables 4 and 5.

**Supplemental Table 1.** List of MEME motifs predicted from ZAR1 in angiosperms.

**Supplemental Table 2.** List of MEME motifs predicted from ZAR1-SUB.

**Supplemental Table 3.** Comparison of MEME motifs between ZAR1-SUB and ZAR1.

**Supplemental Table 4.** List of MEME motifs predicted from ZAR1-CIN.

**Supplemental Table 5.** Comparison of MEME motifs between ZAR1-CIN and ZAR1.

**Supplemental Data Set 1.** List of ZAR1 in angiosperms.

**Supplemental Data Set 2.** List of plant species with the number of ZAR1, ZAR1-SUB and ZAR1-CIN genes.

**Supplemental Data Set 3.** List of the closest NLR genes to ZAR1 locus.

**Supplemental Data Set 4.** Genome loci of ZAR1 and ZRK genes.

**Supplemental Data Set 5.** Amino acid sequences of full-length ZRKs.

**Supplemental Data Set 6.** Amino acid alignment file of 35 ZRK in angiosperms.

**Supplemental Data Set 7.** List of genes genetically linked to ZAR1 in eudicots.

**Supplemental Data Set 8.** List of ZAR1-SUB.

**Supplemental Data Set 9.** List of ZAR1-CIN.

**Supplemental Data Set 10.** Amino acid alignment file of 120 ZAR1 in angiosperms.

**Supplemental Data Set 11.** Reference genome databases used for NLR annotation with NLR-parser.

**Supplemental Data Set 12.** Amino acid sequences of full-length NLRs used for phylogenetic analysis in Figure 1B.

**Supplemental Data Set 13.** Amino acid sequences of full-length NLRs used for phylogenetic analysis in Supplemental Figure 1.

**Supplemental Data Set 14.** Amino acid sequences of 120 ZAR1 in angiosperms.

**Supplemental Data Set 15.** Amino acid sequences of 129 ZAR1-SUB. **Supplemental Data Set 16.** Amino acid sequences of 11 ZAR1-CIN.

**Supplemental Data Set 17.** NLR phylogenetic tree file in Figure 1B.

**Supplemental Data Set 18.** NLR phylogenetic tree file in Supplemental Figure 1.

**Supplemental Data Set 19.** NLR phylogenetic tree file in Figure 2.

**Supplemental Data Set 20.** ZRK phylogenetic tree file in Figure 5A.

**Supplemental Data Set 21.** NLR phylogenetic tree file in Figure 8.

**Supplemental Data Set 22.** Amino acid alignment file of 129 ZAR1-SUB.

**Supplemental Data Set 23.** Amino acid alignment file of 11 ZAR1-CIN.

**Supplemental Data Set 24.** The ConSurf conservation score among ZAR1 proteins.

**Supplemental Data Set 25.** The ConSurf conservation score among ZAR1-SUB proteins.

**Supplemental Data Set 26.** The ConSurf conservation score among ZAR1-CIN proteins.

**Supplemental Data Set 27.** Plasmid list used in this study.

## ACKNOWLEDGEMENTS

We are thankful to several colleagues for discussions and ideas. We thank Sebastian Schornack (Sainsbury Laboratory, University of Cambridge, Cambridge, UK) for valuable comments on this paper. This work was funded by the Gatsby Charitable Foundation, Biotechnology and Biological Sciences Research Council (BBSRC, UK), and European Research Council (ERC BLASTOFF projects). JLGH acknowledges support from the South Dakota Agricultural Experiment Station (SD00H605-16). We thank the Prime Minister of the United Kingdom for announcing a stay-at-home order on 23rd March 2020.

## AUTHOR CONTRIBUTIONS

Conceptualization: H.A., S.K.; Data curation: H.A., T.S., J.K., H.P., J.L.G.H., A.M.; Formal analysis: H.A., T.S., A.M.; Investigation: H.A., T.S., A.M.; Methodology: H.A., T.S., J.K., A.M.; Resources: H.A., T.S., J.K., H.P.; Supervision: H.A., A.M., S.K.; Funding acquisition: S.K.; Project administration: S.K.; Writing initial draft: H.A., J.K., S.K.; Editing: H.A., T.S., J.K., J.L.G.H., S.K.

