## Supplementary material for "Jurassic NLR: conserved and dynamic evolutionary features of the atypically ancient immune receptor ZAR1": SupplementalTable_1_ZAR1_MEME.pdf

**Supplemental Table 1. List of MEME motifs predicted from ZAR1 in angiosperms.**

| Motif ID | Motif logo | E-value | Start | End | Query ID | Domain | Known motif |
|---|---|---|---|---|---|---|---|
| Motif_1 | 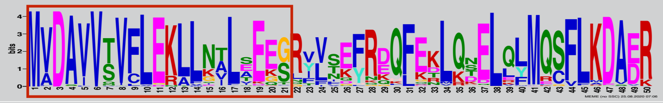 | 6.9e-4272 | 1 | 50 | AtZAR1 | CC | MADA |
| Motif_2 | 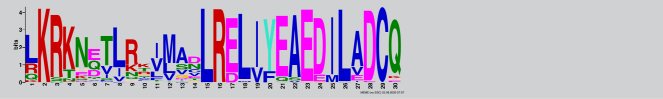 | 1.1e-2467 | 51 | 80 | AtZAR1 | CC | |
| Motif_3 | 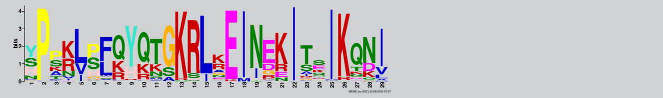 | 6.2e-2106 | 101 | 129 | AtZAR1 | CC | |
| Motif_4 | 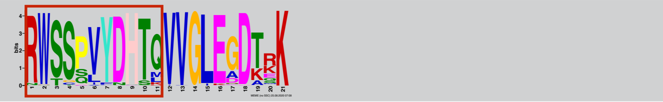 | 3.0e-2096 | 149 | 169 | AtZAR1 | NB-ARC | NBD-NBD interface |
| Motif_5 | 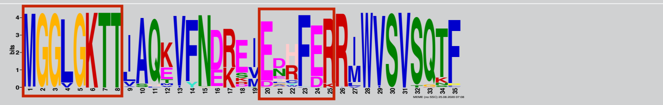 | 1.1e-3624 | 190 | 224 | AtZAR1 | NB-ARC | P-loop, underside surface |
| Motif_6 | 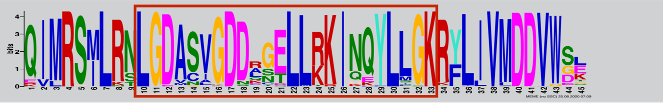 | 5.4e-4559 | 228 | 272 | AtZAR1 | NB-ARC | Underside surface |
| Motif_7 | 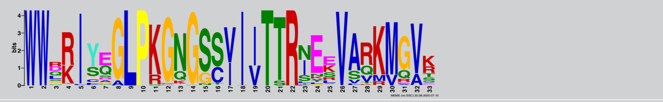 | 2.9e-2989 | 276 | 308 | AtZAR1 | NB-ARC | |
| Motif_8 | 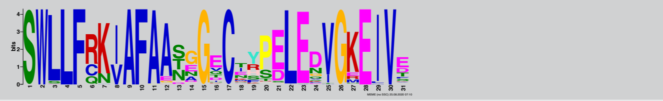 | 1.1e-3059 | 323 | 353 | AtZAR1 | NB-ARC | |
| Motif_9 | 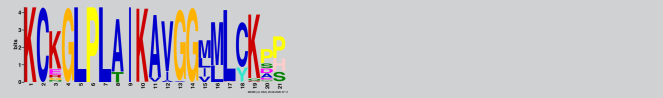 | 6.5e-2078 | 354 | 374 | AtZAR1 | NB-ARC | |
| Motif_10 | 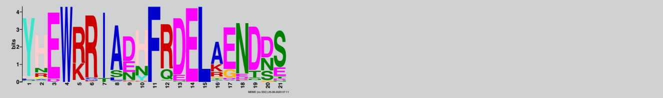 | 2.0e-1941 | 376 | 396 | AtZAR1 | NB-ARC | |
| Motif_11 | 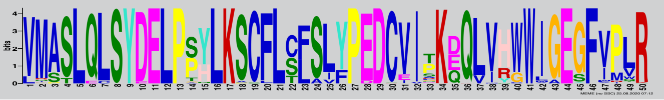 | 1.7e-5508 | 400 | 449 | AtZAR1 | NB-ARC | |
| Motif_12 | 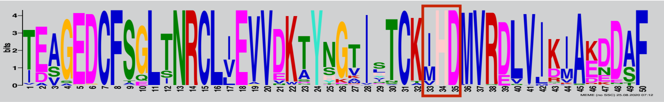 | 3.3e-5399 | 455 | 504 | AtZAR1 | NB-ARC | MHD |
| Motif_13 | 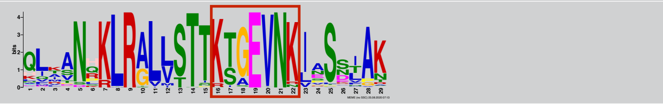 | 2.7e-2246 | 525 | 553 | AtZAR1 | LRR | RLCK interface |
| Motif_14 | 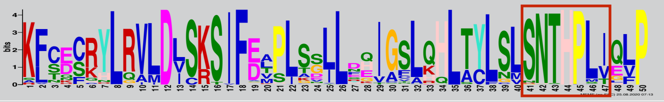 | 5.7e-4432 | 554 | 603 | AtZAR1 | LRR | RLCK interface |
| Motif_15 | 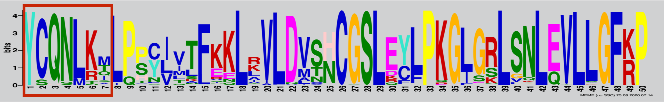 | 2.8e-4683 | 619 | 668 | AtZAR1 | LRR | RLCK interface |
| Motif_16 | 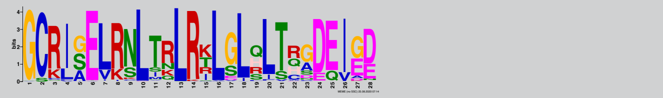 | 1.2e-2177 | 674 | 701 | AtZAR1 | LRR | |
| Motif_17 | 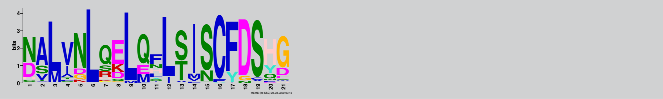 | 3.6e-1546 | 705 | 725 | AtZAR1 | LRR | |
| Motif_18 | 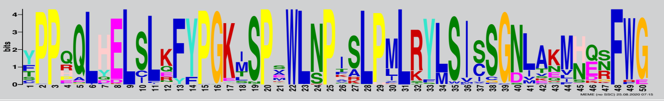 | 3.8e-4856 | 736 | 785 | AtZAR1 | LRR | |
| Motif_19 | 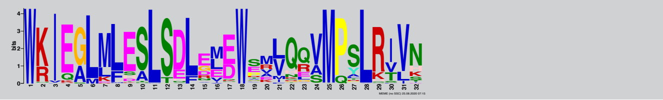 | 6.8e-2538 | 791 | 822 | AtZAR1 | LRR | |
| Motif_20 | 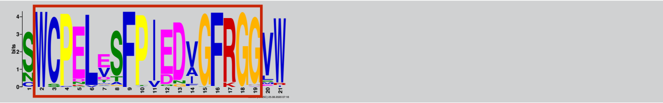 | 4.7e-2129 | 824 | 844 | AtZAR1 | LRR | RLCK interface |
