## Supplementary material for "Jurassic NLR: conserved and dynamic evolutionary features of the atypically ancient immune receptor ZAR1": SupplementalTable_2_ZAR1_SUB_MEME.pdf

**Supplemental Table 2. List of MEME motifs predicted from ZAR1-SUB.**

| Motif ID | Motif logo | E-value | Start | End | Query ID | Domain | Known motif |
|---|---|---|---|---|---|---|---|
| Motif_1 | 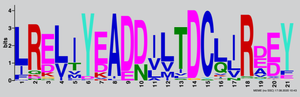 | 1.2e-1593 | 69 | 89 | XP_004243429.1 | CC | |
| Motif_2 | 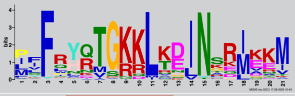 | 1.5e-1173 | 105 | 125 | XP_004243429.1 | CC | |
| Motif_3 | 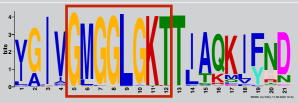 | 2.5e-1939 | 192 | 212 | XP_004243429.1 | NB-ARC | P-loop |
| Motif_4 | 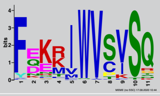 | 8.0e-701 | 219 | 229 | XP_004243429.1 | NB-ARC | |
| Motif_5 | 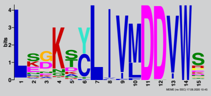 | 1.6e-1072 | 263 | 277 | XP_004243429.1 | NB-ARC | |
| Motif_6 | 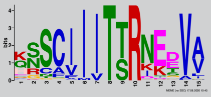 | 1.6e-1022 | 296 | 310 | XP_004243429.1 | NB-ARC | |
| Motif_7 | 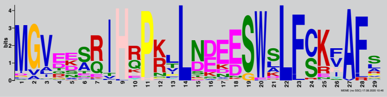 | 4.0e-2073 | 313 | 341 | XP_004243429.1 | NB-ARC | |
| Motif_8 | 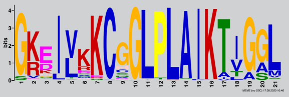 | 1.2e-1683 | 356 | 376 | XP_004243429.1 | NB-ARC | |
| Motif_9 | 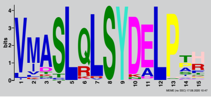 | 1.8e-1099 | 408 | 422 | XP_004243429.1 | NB-ARC | |
| Motif_10 | 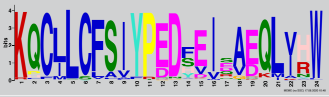 | 6.5e-2273 | 424 | 447 | XP_004243429.1 | NB-ARC | |
| Motif_11 | 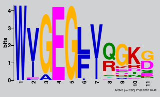 | 5.2e-677 | 448 | 458 | XP_004243429.1 | NB-ARC | |
| Motif_12 | 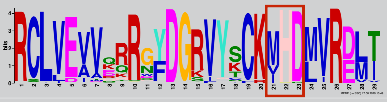 | 1.7e-2710 | 476 | 504 | XP_004243429.1 | NB-ARC | MHD |
| Motif_13 | 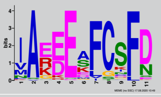 | 4.5e-686 | 507 | 517 | XP_004243429.1 | NB-ARC | |
| Motif_14 | 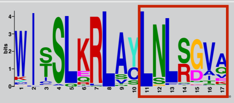 | 1.1e-1225 | 597 | 613 | XP_004243429.1 | LRR | RLCK interface |
| Motif_15 | 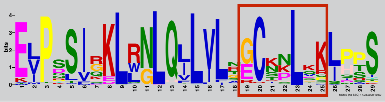 | 6.5e-1849 | 617 | 645 | XP_004243429.1 | LRR | RLCK interface |
| Motif_16 | 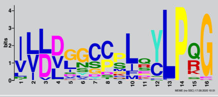 | 4.9e-1104 | 654 | 669 | XP_004243429.1 | LRR | |
| Motif_17 | 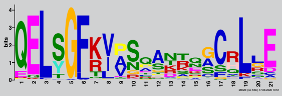 | 2.3e-1036 | 677 | 697 | XP_004243429.1 | LRR | |
| Motif_18 | 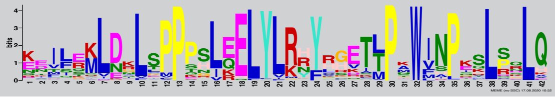 | 4.8e-3168 | 744 | 785 | XP_004243429.1 | LRR | |
| Motif_19 | 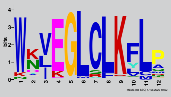 | 1.4e-716 | 810 | 821 | XP_004243429.1 | LRR | |
| Motif_20 | 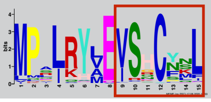 | 3.1e-927 | 834 | 848 | XP_004243429.1 | LRR | RLCK interface |
