## Supplementary material for "Jurassic NLR: conserved and dynamic evolutionary features of the atypically ancient immune receptor ZAR1": SupplementalTable_4_ZAR1_CIN_MEME.pdf

**Supplemental Table 4. List of MEME motifs predicted from ZAR1-CIN.**

| Motif ID | Motif logo | E-value | Start | End | Query ID | Domain | Known motif |
|---|---|---|---|---|---|---|---|
| Motif_1 | 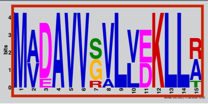 | 4.1E-28 | 1 | 15 | RWR856 56.1 | CC | MADA |
| Motif_2 | 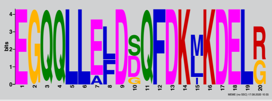 | 1.7E-61 | 20 | 39 | RWR856 56.1 | CC | |
| Motif_3 | 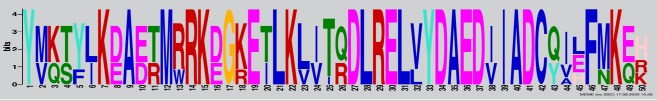 | 1.2E-222 | 40 | 89 | RWR856 56.1 | CC | |
| Motif_4 | 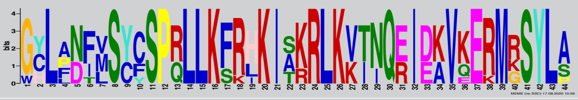 | 1.5E-171 | 91 | 134 | RWR856 56.1 | CC | |
| Motif_5 | 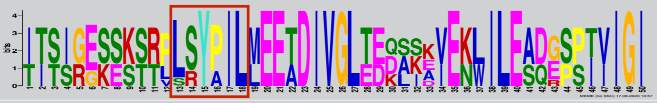 | 3.7E-155 | 139 | 188 | RWR856 56.1 | NB-ARC | NBD-NBD interface |
| Motif_6 | 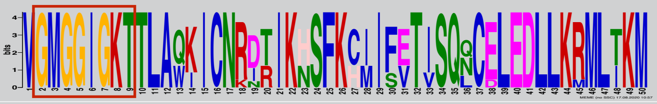 | 7.6E-256 | 189 | 238 | RWR856 56.1 | NB-ARC | P-loop |
| Motif_7 | 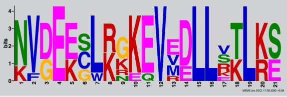 | 7.2E-33 | 239 | 259 | RWR856 56.1 | NB-ARC | |
| Motif_8 | 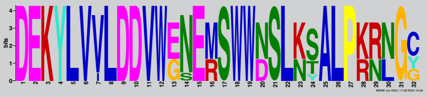 | 8.0E-150 | 263 | 294 | RWR856 56.1 | NB-ARC | |
| Motif_9 | 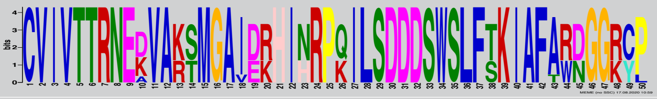 | 7.2E-275 | 295 | 344 | RWR856 56.1 | NB-ARC | |
| Motif_10 | 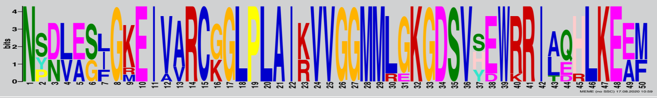 | 2.6E-224 | 345 | 394 | RWR856 56.1 | NB-ARC | |
| Motif_11 | 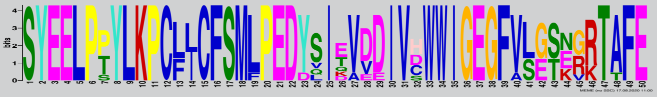 | 3.3E-249 | 410 | 459 | RWR856 56.1 | NB-ARC | |
| Motif_12 | 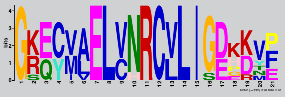 | 7.7E-57 | 461 | 481 | RWR856 56.1 | NB-ARC | |
| Motif_13 | 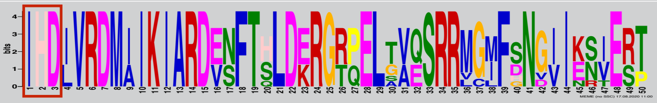 | 2.0E-216 | 490 | 539 | RWR856 56.1 | NB-ARC | MHD |
| Motif_14 | 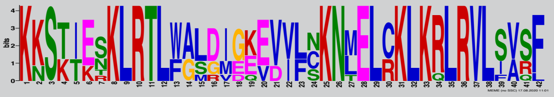 | 7.2E-128 | 542 | 583 | RWR856 56.1 | LRR | |
| Motif_15 | 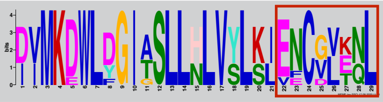 | 1.1E-98 | 591 | 619 | RWR856 56.1 | LRR | RLCK interface |
| Motif_16 | 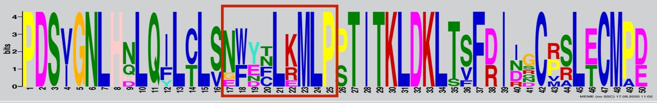 | 6.9E-219 | 620 | 669 | RWR856 56.1 | LRR | RLCK interface |
| Motif_17 | 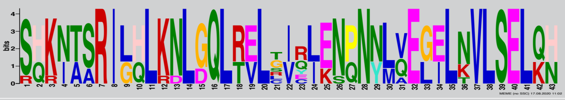 | 1.3E-143 | 689 | 731 | RWR856 56.1 | LRR | |
| Motif_18 | 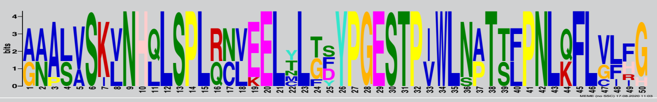 | 6.1E-173 | 747 | 796 | RWR856 56.1 | LRR | |
| Motif_19 | 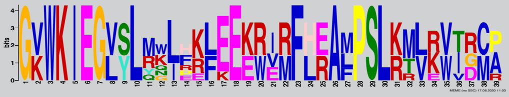 | 8.2E-93 | 812 | 850 | RWR856 56.1 | LRR | |
| Motif_20 | 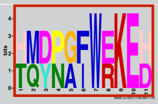 | 1.7E-29 | 860 | 870 | RWR856 56.1 | LRR | RLCK interface |
