## Supplementary material for "Jurassic NLR: conserved and dynamic evolutionary features of the atypically ancient immune receptor ZAR1": SupplementalTable_5_ZAR1_CIN_MEME_Compare.pdf

### Supplemental Table 5. Comparison of MEME motifs between ZAR1-CIN and ZAR1.

| ZAR1-CIN motif ID | Motif logo | ZAR1 motif ID | Motif logo | Domain |
|---|---|---|---|---|
| Motif_1 | 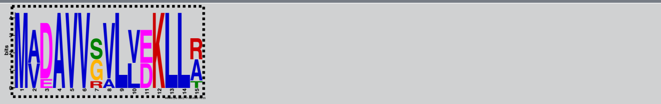 | Motif_1 | 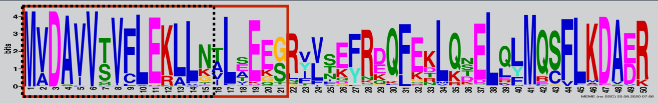 | CC |
| Motif_2 | 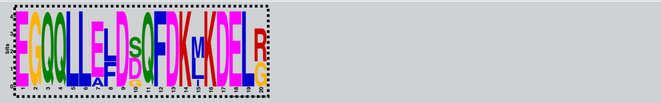 | Motif_1 | 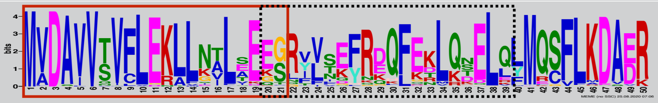 | CC |
| Motif_3 | 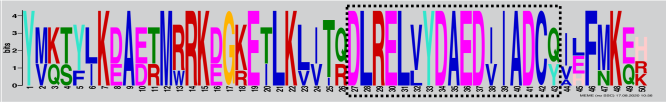 | Motif_2 | 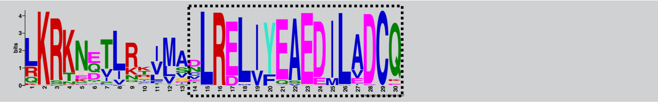 | CC |
| Motif_4 | 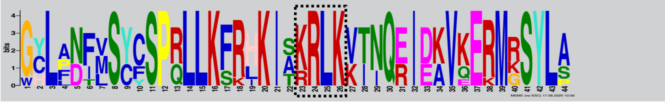 | Motif_3 | 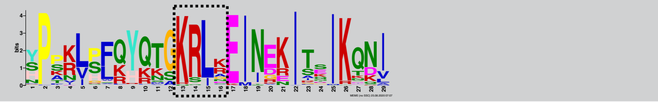 | CC |
| Motif_5 | 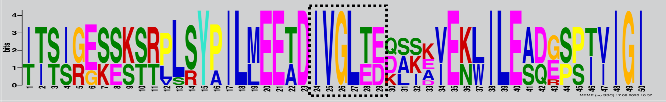 | Motif_4 | 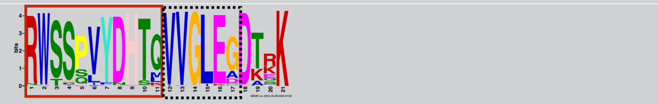 | NB-ARC |
| Motif_6 | 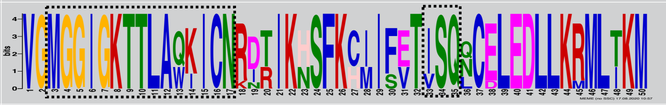 | Motif_5 | 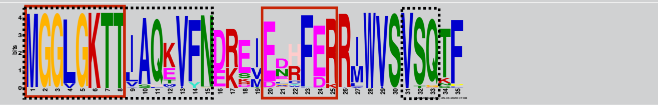 | NB-ARC |
| Motif_7 | 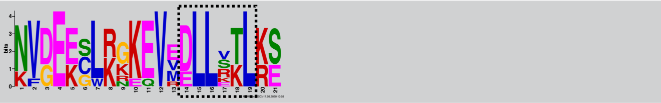 | Motif_6 | 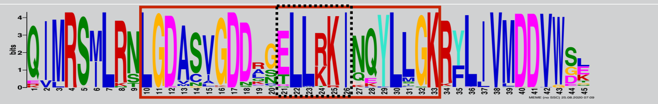 | NB-ARC |
| Motif_8 | 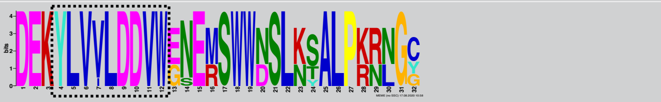 | Motif_6 | 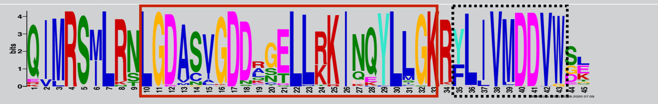 | NB-ARC |
| Motif_9 | 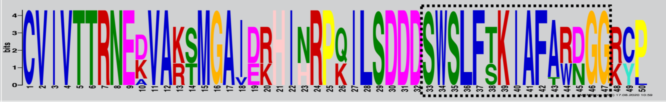 | Motif_8 | 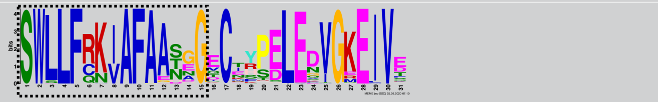 | NB-ARC |
| Motif_10 | 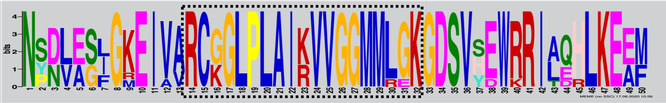 | Motif_9 | 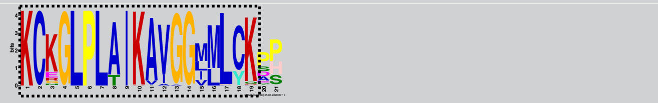 | NB-ARC |
| Motif_11 | 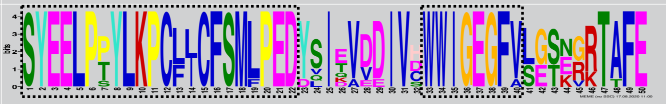 | Motif_11 | 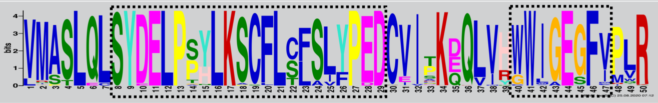 | NB-ARC |
| Motif_12 | 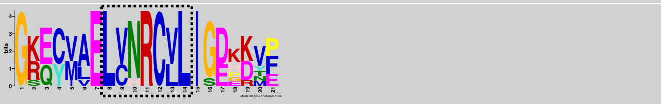 | Motif_12 | 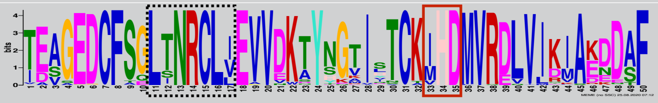 | NB-ARC |
| Motif_13 | 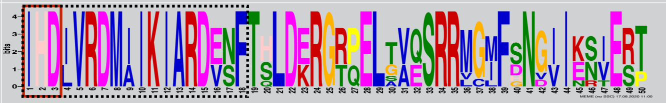 | Motif_12 | 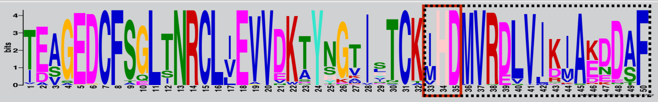 | NB-ARC |
| Motif_14 | 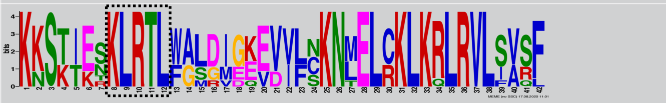 | Motif_13 | 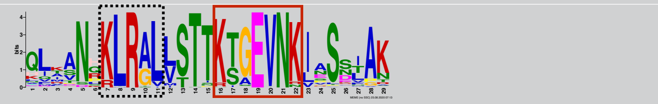 | LRR |
| Motif_15 | 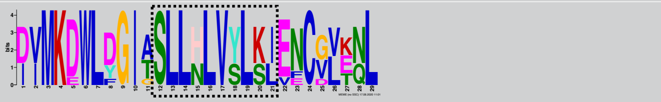 | Motif_14 | 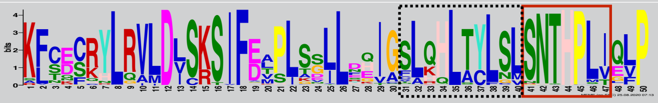 | LRR |
| Motif_16 | 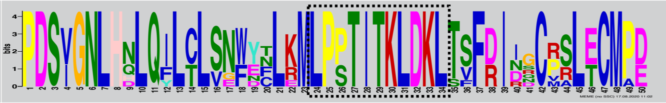 | Motif_15 | 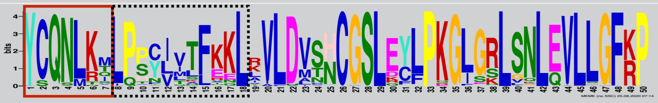 | LRR |
| Motif_17 | 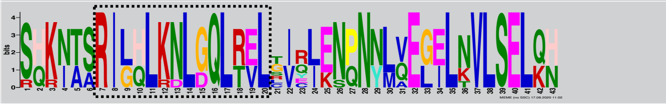 | Motif_16 | 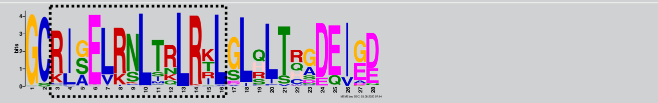 | LRR |
| Motif_18 | 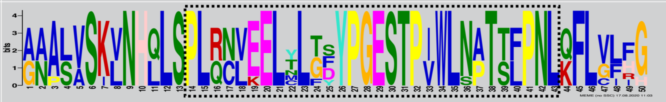 | Motif_18 | 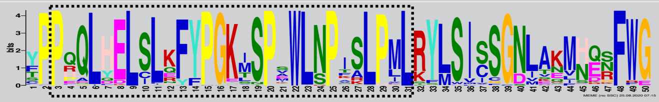 | LRR |
| Motif_19 | 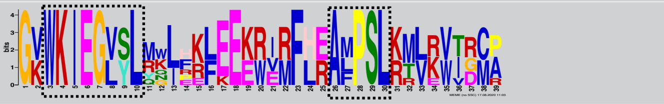 | Motif_19 | 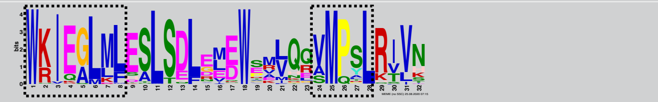 | LRR |
| Motif_20 | 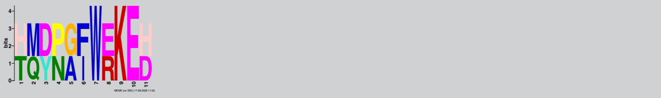 | ND | | LRR |

*Black dot boxes indicate corresponding regions between ZAR1-CIN and ZAR1 based on MAFFT v7 alignment in Supplemental Figure 15. Red boxes indicate motifs which are highlighted in Figure 3 .
